# FGF independent MEK1/2 signalling is essential for male fetal germline development in mice

**DOI:** 10.1101/2023.02.05.527224

**Authors:** Rheannon O. Blücher, Rachel S. Lim, Ellen G. Jarred, Matthew E. Ritchie, Patrick S. Western

## Abstract

**Background:** Germline development provides the founding cells for spermatogenesis and oogenesis in males and females, respectively. Disrupted germline differentiation or compromised testis development can lead to subfertility or infertility and are strongly associated with testis cancer in humans. In mice, SRY and SOX9 induce expression of a range of genes, including *Fgf9,* that promote Sertoli cell differentiation and testis development. FGF9 is also thought to promote male germline differentiation but the pathway through which it signals is unknown. As FGFs signal through Mitogen-Activated Protein Kinases (MAPKs) in other tissues, we explored whether FGF9 regulates male germline development through MAPK by inhibiting either FGF or MEK1/2 signalling in fetal testis cultures from embryonic day (E)12.5, immediately after gonadal sex determination and testis cord formation, but prior to male germline commitment.

**Results:** Inhibition of MEK1/2 disrupted mitotic arrest, dysregulated a broad range of male germline development genes and prevented the upregulation of key male germline markers DPPA4 and DNMT3L. In contrast, when FGF signalling was inhibited, the male germline specific transcriptional program and the expression of male germline markers DPPA4 and DNMT3L were unaffected, and germ cells entered mitotic arrest normally. While male germline development was not disrupted by FGF inhibition, some genes were commonly altered after 24h of FGF or MEK1/2 inhibition including genes involved in maintenance, germline stem cells, Nodal signalling, proliferation, and germline cancer.

**Conclusions:** Together, these data demonstrate a novel and essential role for MEK1/2 signalling in male germline differentiation, but a more limited role for FGF signalling. Our data strongly indicate that additional ligands act through MEK1/2 to promote male germline differentiation and highlight a need for further mechanistic understanding of male germline development.

## Introduction

Development of functional sperm and oocytes is essential for fertility and transmission of genetic and epigenetic information to offspring. A critical part of germline development involves the commitment of germ cells to male or female development in response to somatic cell signalling (Adams and McLaren, 2002). Germ cells within a developing testis commit to male development from embryonic day (E)12.5 and enter mitotic arrest between E13.5 and E15.5 (Adams and McLaren, 2002, Western et al., 2008). They then re-enter the cell cycle and establish spermatogonial stem cells (SSCs) before entering meiosis during spermatogenesis in post-natal life. In contrast, commitment to female germline in the ovary is closely followed by entry into meiosis by E15.5 (Western et al., 2008, Adams and McLaren, 2002, Miles et al., 2010).

Male germline differentiation depends on development of an appropriate testicular environment. In XY mice, *Sry* (Sex region, Y-chromosome) and SOX9 (SRY box gene 9) drive pre-supporting cell commitment to Sertoli cell development at E11.5 (Koopman et al., 1991, Kent et al., 1996). Sertoli cells form testis cords that enclose germ cells, a process supported by FGF9 (fibroblast growth factor 9), which further induces *Sox9* expression and drives Sertoli cell proliferation in conjunction with prostaglandin D_2_ (PGD2) (Kim et al., 2006, Colvin et al., 2001, Adams and McLaren, 2002, Wilhelm et al., 2005). *Fgf9* expression peaks at E11.5, before declining to low levels by E13.5 (Bowles et al., 2010, Colvin et al., 2001, Kim et al., 2006, Yildirim et al., 2020, Jameson et al., 2012b). Loss of function mutations in *Fgf9* or its receptor, *Fgfr2*, disrupt testis development leading to ovary or ovotestis formation, reduced germ cell numbers and germline sex-reversal (DiNapoli et al., 2006, Kim et al., 2006, Colvin et al., 2001, Bagheri-Fam et al., 2008, Bagheri-Fam et al., 2017).

Evidence suggests that FGF9 also directly promotes male germline development by acting as a meiosis inhibiting factor by repressing *Stra8* (Stimulated by retinoic acid gene 8) and ensuring expression of male germline genes including *Nanos2* and *Dnmt3l* (Bowles et al., 2010, Barrios et al., 2010). However, male germ cells develop in *Wnt4/Fgf9* double null mice (Jameson et al., 2012a), indicating that *Fgf9* is dispensable in this double null background. Furthermore, the mechanism through which FGF9 promotes male germ cell development in wild type mice remains unknown.

FGF ligands, including FGF9, are also required for the establishment of pluripotent embryonic germ cells (EGCs) from primordial germ cells (PGCs) in culture and promote proliferation and an undifferentiated state in SSCs (Durcova-Hills et al., 2006, Kanatsu-Shinohara et al., 2014, Yang et al., 2021). Consistent with this, in gonads cultures, ectopic FGF9 maintained expression of the pluripotency genes *Sox2* and *Oct4* in XY germ cells (Bowles et al., 2010, Gustin et al., 2016b). Moreover, isolated E11.5 or E12.5 XY germ cells exposed to high levels of FGF9 maintained proliferation, but low levels of FGF9 promoted male germ cell differentiation (Ulu et al., 2017). *Fgf9* is downregulated by E13.5 and the potential for germ cells to make EGCs is lost from E12.5, consistent with germ cell entry into mitotic arrest and repression of pluripotency between E13.5 and E15.5 (Durcova-Hills et al., 2008, Western et al., 2010, Bowles et al., 2010).

FGF ligands bind FGF receptors and rapidly activate intracellular responses via MAP kinase signalling, including through MEK1/2 and phosphorylation of its target ERK1/2 (Dorey and Amaya, 2010, Weyman and Wolfman, 1998, Lovicu and McAvoy, 2001). Low levels of ERK1/2 phosphorylation were detected in isolated E11.5 XX and XY germ cells, but gradually increased between E12.5 and E14.5 (Sorrenti et al., 2020), highlighting a potential role for MEK1/2 signalling during fetal germ cell development. Moreover, inhibiting ERK1/2 signalling maintains stem cells in ground state pluripotency (Ying et al., 2008), demonstrating a role for ERK1/2 in priming differentiation of pluripotent cells, an activity facilitated by the E26 transformation-specific (ETS) factors, ETV4 and ETV5 (Kalkan et al., 2019, Mulas et al., 2019).

A caveat of previous studies that have deleted *Fgf9* or *Fgfr2* is that germline developmental outcomes are potentially confounded by somatic-sex reversal mediated by loss of FGF signalling. We hypothesised that FGF9 signalling through MEK1/2 promotes male germline development soon after supporting cells commit to Sertoli cell development and testis differentiation. We inhibited FGF receptor or MEK1/2 signalling at E12.5, after commitment to testis development at E11.5 but before male germline differentiation and entry to mitotic arrest. Our data demonstrate that inhibiting MEK1/2 signalling at E12.5 allows germ cells to escape mitotic arrest and substantially disrupts male germline differentiation. This indicates that XY germ cells require FGF-independent MEK1/2 signalling to successfully mediate male germline differentiation.

## Results

### Inhibition of FGF or MEK1/2 signalling disrupts testis development

To determine the role of FGF and potential downstream target signalling pathways required for male germ cell development, E12.5 XY gonad-mesonephros samples were cultured with vehicle control (DMSO) or a range of small molecule inhibitors of FGF receptors (FGFR1-3), MEK1/2, p38MAPK and PI3K. These included pan-FGFR (FGFR1-3) inhibitor BGJ398/Infigratinib (FGFRi; IC50 ∼1.0nM), MEK1/2 inhibitor PD0325901/Mirdametinib (MEKi; IC50 0.33nM), p38MAPK inhibitor PH-797804 (p38i; IC50 26nM) or PI3K inhibitor GSK1059615 (PI3Ki, IC50 5nM; Table 1). Importantly all of these inhibitors have IC50 values in the low nM range and have been extensively validated in clinical trials demonstrating their high specificity, potency and cell tolerance (Table 1).

**Table 1:** Summary of treatments and doses used in gonad cultures.

| Treatment | Dose | IC50 and target | Clinical trial phase, clinicaltrials.gov identifier | Supplier, catalogue number | References |
| --- | --- | --- | --- | --- | --- |
| DMSO (vehicle control) | Equal to DMSO in drug treatments ($\geq 1/5000$) | NA | NA | Thermo-Fisher, D12345 | |
| BGJ398/ Infigratinib (FGFRi) | 125, 250, 500, 1000 and 2500 nM | IC50 – FGFR1-3: 0.9-1.4 nM, FGFR4: 60 nM | Phase II, NCT02150967 | SelleckChem, S2183 | (Javle et al., 2018) |
| PD0325901/ Mirdametinib (MEKi) | 125, 250, 500 and 1000 nM | IC50 – MEK: 0.33 nM | Phase II, NCT03962543 | SelleckChem or MedChem Express, S1036 or HY-10254 | (Weiss et al., 2021) |
| PH-797804 (p38i) | 500 nM | IC50 – p38α: 26 nM; p38β 102 nM | Phase II, NCT00559910 | MedChem Express, HY-10403 | (MacNee et al., 2013) |
| GSK1059615 (PI3Ki) | 500 nM | IC50 – PI3Kα/ $\beta$/$\delta$/$\gamma$: 0.4-5 nM, mTOR: 12 nM | Phase I, NCT00695448 | MedChem Express, HY-12036 | |
| Recombinant mouse FGF9 | 50 ng/mL | FGFR1-3 | NA | R&D systems, 7399-F9-025 | (Gustin et al., 2016b). |
| AZD4547 (second FGFR inhibitor) | 125, 250 and 500 nM | IC50 – FGFR1-3: 0.2-2.5 nM; FGFR4: 165 nM; VEGFR2: 24 nM | Phase II, NCT02465060 | SelleckChem, S2801 | (Chae et al., 2020) |
| SU5402 (third FGFR inhibitor) | 5000nM | IC50 VEGFR: 20nM; FGFR1: 30nM | N/A | MedChem Express, HY-10407 | (Sun et al., 1999) |
| Ralimetinib dimesylate/ LY2228820 (second p38 inhibitor) | 500 nM | IC50 – p38α: 5.3 nM; p38β: 5.3 nM | Phase I, NCT01663857 | MedChem Express, HY-13241 | (Patnaik et al., 2016) |
| PF-04691502 (second PI3K inhibitor) | 500 nM | IC50 – PI3Kα/ $\beta$/$\delta$/$\gamma$: 1.6-2.1 nM; mTOR: 16 nM | Phase II, NCT01420081 | MedChem Express, HY-15177 | (del Campo et al., 2016) |

To determine whether FGFR, MEK1/2, p38MAPK or PI3K disrupted germ cell mitotic arrest, we initially used a concentration of 500nM based on the low IC50 values for each drug and our observations that drugs with similar IC50s maximally inhibit their targets in gonad cultures in the 100-1000nM range, with the difference between cell free IC50 values and efficacy in culture presumably due to drug bioavailability in cells and whole organs. The vehicle control, DMSO, was used at a dilution of ≥1/5,000 in all experiments, a concentration that does not affect gonad or germline development (Miles et al., 2013, Gustin et al., 2016b). Bright-field and fluorescence examination of E12.5 testis-mesonephros samples cultured with control (DMSO) or drug for 72h, provided an initial readout of the impact of each drug based on germ cell organisation within testis cords, marked by germ cell specific expression of *Oct4-GFP* (Fig 1A). DMSO controls developed well-defined cords containing germ cells, but FGFRi and MEKi treatment resulted in poor testis cord structure and GFP positive germ cells scattered throughout the testis. p38i and PI3Ki treated gonads were morphologically similar to DMSO controls, with GFP positive germ cells contained within well-defined testis cords (Fig 1A).

**Figure 1:**
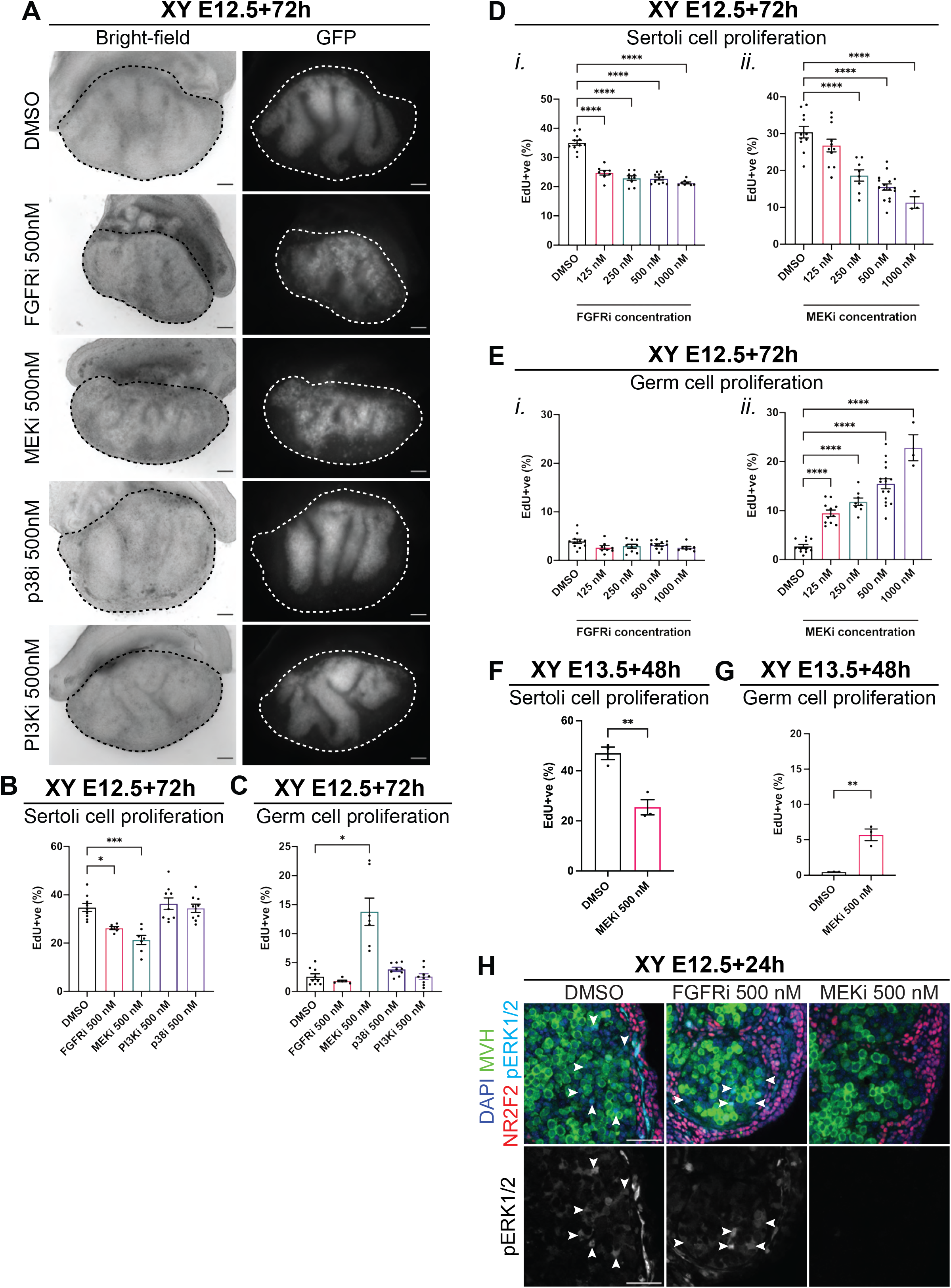
FGF and MEK1/2 inhibition disrupts fetal testis development but only MEK1/2 inhibition disrupts germ cell mitotic arrest. **A)** Bright-field and GFP images of E12.5 XY gonad-mesonephros tissue cultured for 72h with DMSO, or 500nM of FGFRi, MEKi, p38i or PI3Ki. Scale bar: 100μm. Dotted lines highlight the gonad. **B-E)** Flow cytometric analysis of Sertoli (B and D) or germ (C and E) cell proliferation based on EdU incorporation in XY E12.5 gonad-mesonephros tissue cultured for 72h with DMSO, 500nM of FGFRi, MEKi, PI3Ki or p38i (B-C) or 125, 250, 500 or 1000nM of FGFRi or MEKi (D-E). **F-G)** Flow cytometric analysis of Sertoli (F) or germ (G) cell proliferation based on EdU incorporation in XY E13.5 gonad-mesonephros tissue cultured for 48h with DMSO or 500nM of FGFRi or MEKi. **H)** Immunofluorescent images of E12.5 gonad-mesonephros tissue cultured for 24h with DMSO, 500nM of FGFRi or MEKi demonstrating MEK1/2 signalling activity. Top panel: DAPI (blue), MVH (green), NR2F2 (red), pERK1/2 (cyan). Bottom panel: pERK1/2 (grey). Scalebar represents 50μm. Replicates: A-C) n=6-9, D-E) n=3-16, F-G) n=3, H) n=3-4. Statistics: B, D-E) Ordinary one-way ANOVA with Tukey’s multiple comparison, C) Brown-Forsythe and Welch ANOVA with Dunnett’s T3 multiple comparisons, F-G) Unpaired two-tailed t-test. Error bars: Mean ± SEM. Significance between control and treatment: *P<0.05, **P<0.01, ***P<0.001, ****P<0.0001.

To further determine the effects of each treatment on testis development, we added 5-ethynyl-2’-deoxyuridine (Edu) during the final 2h of culture and used flow cytometry to assess proliferation based on EdU incorporation during S-phase and DNA content using propidium iodide staining in E12.5 gonad-mesonephric samples cultured for 72h with each inhibitor (Miles et al., 2013, Gustin et al., 2016b, Gustin et al., 2016a) (Fig 1B-C, S1). As expected, Sertoli cells were highly proliferative in DMSO controls (Fig 1B). However, FGFR or MEK1/2 inhibition reduced Sertoli cell proliferation compared to XY DMSO controls (*P*<0.05 or *P*<0.001, respectively, Fig 1B). As expected, germ cell proliferation was very low in DMSO controls (Fig 1C), confirming that the germ cells had entered mitotic arrest, a key milestone in male germline differentiation (Western et al., 2008). In contrast, germ cell proliferation was substantially higher in MEKi treated gonads (*P*<0.05), demonstrating that germ cell mitotic arrest was disrupted (Fig 1C). However, the percentage of germ cells incorporating EdU in FGFRi-treated testes was similar to DMSO controls, demonstrating that the germ cells had entered mitotic arrest (Fig 1C). As FGFRi was expected to disrupt germ cell mitotic arrest, this outcome was confirmed using another potent FGFR inhibitor (AZD4547; Table 1), which also resulted in reduced Sertoli cell proliferation but did not disrupt germ cell mitotic arrest (Fig S2A-B). Neither p38i (PH-797804) nor PI3Ki (GSK1059615) affected Sertoli cell proliferation or germ cell mitotic arrest (Fig 1B-C), an outcome confirmed using independent, p38 and PI3K inhibitors, ralimetanib dimesylate (LY2228820) and PF-04691502 (Table 1; Fig S2C-D).

Given both FGFRi and MEKi reduced Sertoli cell proliferation and disrupted testis cord development, we analysed AMH, SOX9 and FOXL2 using immunofluorescence (IF) to ensure that FGFR or MEK inhibition did not result in somatic sex-reversal. Notably, robust SOX9 and AMH staining was detected by IF in male DMSO, FGFRi and MEKi samples but not in female gonads (Fig S3A), indicating that treated XY gonads maintained a male phenotype. Consistent with this, assessment of SOX9 intensity using flow cytometry demonstrated that SOX9 expression was not reduced in FGFRi or MEKi treated samples compared to DMSO controls. (Fig S3B). Furthermore, IF staining for the female marker, FOXL2, revealed strong expression in female gonads, but minimal staining in male DMSO, FGFRi or MEKi samples, although occasional FOXL2 positive cells were detected in MEKi treated samples (Fig S3C). Together these data suggest that FGFRi or MEKi did not result in somatic sex-reversal of the gonads.

### FGF and MEK1/2 signalling are both required for Sertoli cell proliferation, but only MEK1/2 is required for germ cell mitotic arrest

To determine the dose response to MEK1/2 and FGFR inhibition, E12.5 XY gonad-mesonephros samples were cultured with MEKi or FGFRi at 0 (DMSO diluted *≥*1/5,000), 125, 250, 500 and 1000nM and assessed using flow cytometry (Fig S4). Compared to DMSO control, proliferation of SOX9 expressing Sertoli cells was reduced by all doses of FGFRi ≥125nM (*P*<0.0001; Fig 1Di) and ≥250nM MEKi (*P*<0.0001, Fig 1Dii). Of interest, although the maximal impact of FGFRi on Sertoli cell proliferation occurred at 125nM, it did not further reduce Sertoli cell proliferation even at 1000nM and this effect was noticeably less than that of MEKi at doses of 500nM and 1000nM (Fig 1Di vs ii; Fig S2I).

Consistent with our initial observations (Fig 1C), although Sertoli cell proliferation was reduced, germ cell mitotic arrest remained unaffected by FGFRi, even with doses of 1000nM (Fig 1Ei) or 2500nM (Fig S2E-F). SU5402 was previously used at 5000nM to inhibit FGFR (Bowles et al., 2010). Confirming the outcomes obtained using FGFRi and AZD4547, 5000nM SU5402 reduced Sertoli cell proliferation to a similar extent as FGFRi and AZD4547, but it did not affect germ cell mitotic arrest (Table 1; Fig S2E-F). In contrast, MEKi potently disrupted germ cell mitotic arrest even when used at 125nM, with increasingly high proportions of EdU positive proliferative germ cells as MEKi concentration increased (*P*<0.0001, Fig 1Eii).

In repeated experiments MEK1/2 inhibition profoundly disrupted germ cells mitotic arrest in E12.5 XY gonads but three different FGFR inhibitors did not, even though FGFs typically elicit a response through MEK1/2-pERK1/2 within 10-15 minutes (Weyman and Wolfman, 1998, Lovicu and McAvoy, 2001). As FGF9 expression peaks at E11.5 (Jameson et al., 2012b), a potential explanation for the inability of FGFRi to disrupt germ cell mitotic arrest could be that FGF9 was inhibited too late in E12.5 cultures. However, as for *Fgf9* or *Fgfr2* genetic deletions (DiNapoli et al., 2006, Colvin et al., 2001, Kim et al., 2006, Bagheri-Fam et al., 2008, Bagheri-Fam et al., 2017), inhibition of FGFR at E11.5 is expected to cause somatic sex-reversal and consequent germ cell sex-reversal that would substantially confound data interpretation. We were therefore unable to test whether earlier FGFR inhibition disrupted male germline development.

To determine if MEK1/2 inhibition affected Sertoli cell proliferation and germ cell mitotic arrest after E13.5, E13.5 XY gonad-mesonephros samples were cultured for 48h with DMSO or 500nM MEKi. MEKi treatment from E13.5 significantly reduced Sertoli (*P*<0.01, Fig 1F) and increased germ cell proliferation (*P*<0.01, Fig 1G). However, the effect of MEKi on germ cell mitotic arrest appeared to be diminished compared to E12.5 (E13.5: EdU 6%, E12.5: 15%, Fig 1Eii,G), indicating that the ability of MEKi to disrupt mitotic arrest decreased between E12.5 and E13.5.

We previously demonstrated that FGF9 induces proliferation of XX somatic cells at rates similar to XY gonads (Gustin et al., 2016b). To confirm that FGFRi and MEKi effectively blocked FGF9 activity, XX E12.5 gonads were cultured for 48h in media containing DMSO, 50ng/mL FGF9, 500nM FGFRi, 500nM MEKi, FGF9+FGFRi or FGF9+MEKi and assessed using flow cytometry. As expected, FGF9 substantially increased the proliferation of XX gonadal somatic cells compared to XX controls (*P*<0.05, *P*<0.0001; Fig S2G). Critically, both FGFRi and MEKi completely neutralised FGF9, with somatic cell proliferation decreased to XX control levels in FGF9 + FGFRi or FGF9 + MEKi treated XX gonads (Fig S2Gi-ii). In contrast, neither p38i nor PI3Ki counteracted FGF9 induced somatic cell proliferation in XX gonads, indicating that neither p38MAPK nor PI3K regulate primary pathways through which FGF drives somatic cell proliferation in developing gonads (Fig SGiii).

### Inhibition of MEK1/2 completely abolished ERK1/2 phosphorylation in the developing testis, but FGFR inhibition did not

FGF activation of MEK1/2 rapidly results in phosphorylation of ERK1/2, and MEK inhibition blocks this activity (Weyman and Wolfman, 1998, Lovicu and McAvoy, 2001). To determine if inhibition of FGF or MEK1/2 signalling prevented phosphorylation of ERK1/2, E12.5 XY gonad-mesonephros were cultured with 500nM FGFRi or MEKi for 24h and phosphorylated ERK1/2 (pERK1/2) was assessed using IF (Fig 1H, S2H). Surprisingly, pERK1/2 was not detected in MVH expressing germ cells in the developing testis. However, consistent with MEK1/2 activity in Sertoli cells, pERK1/2 was detected at low levels in MVH negative somatic cells within testis cords in DMSO controls. While there are no other somatic cells within testis cords, Sertoli cell localisation of pERK1/2 could not be definitively determined using SOX9 IF as the SOX9 and pERK1/2 antibodies were both raised in rabbit. Robust pERK1/2 was also detected in somatic cells outside of the testis cords that appeared be endothelial cells, however, this was not confirmed. pERK1/2 was not detected in MEKi treated samples, although it was detected in Sertoli and somatic cells outside of the testis cords in FGFRi treated gonads (Fig 1H, S2H) demonstrating that MEKi abolished ERK1/2 phosphorylation, but FGFR inhibition did not. Consistent with this, MEKi reduced Sertoli cell proliferation to a greater extent than FGFRi (P<0.0001 MEKi vs FGFRi, Fig S2I). However, as 500nM of FGFRi completely blocked FGF9 induced proliferation in XX somatic cells (Fig S2Gi), the most likely explanation for the inability of FGFRi to completely eliminate pERK1/2 in Sertoli cells is that MEK1/2 may be activated independently of FGFR, perhaps by PGD2 which is known to promote Sertoli cell proliferation and activate MEK1/2 (Adams and McLaren, 2002, Wilhelm et al., 2005, Kuroyanagi et al., 2014).

### FGF and MEK1/2 signalling is required for normal testis cord formation

As gonad wholemount images indicated testis cords were disrupted by FGFRi or MEKi (Fig 1A), we used IF to investigate MVH positive germ cells relative to SOX9-expressing Sertoli cells, SMA-expressing peritubular myoid cells or laminin, which delineate testis cords. In DMSO controls the majority of Sertoli cells were organised in a single layer at the outer edge of the testis cords, with germ cells very rarely found outside the cords (Fig 2). In FGFRi and MEKi treated samples some Sertoli cells localised to the outer edge of testis cords, but gaps were evident between the Sertoli cells, and many Sertoli cells remained dispersed throughout the interior of the testis cords (Fig 2A). Furthermore, germ cells were occasionally present in the gaps between Sertoli cells at the testis cord basement membrane (Fig 2A) and were mis-localised outside testis cords in FGFRi treated cultures, although this was more common in MEKi treated gonads (Fig 2B).

**Figure 2:**
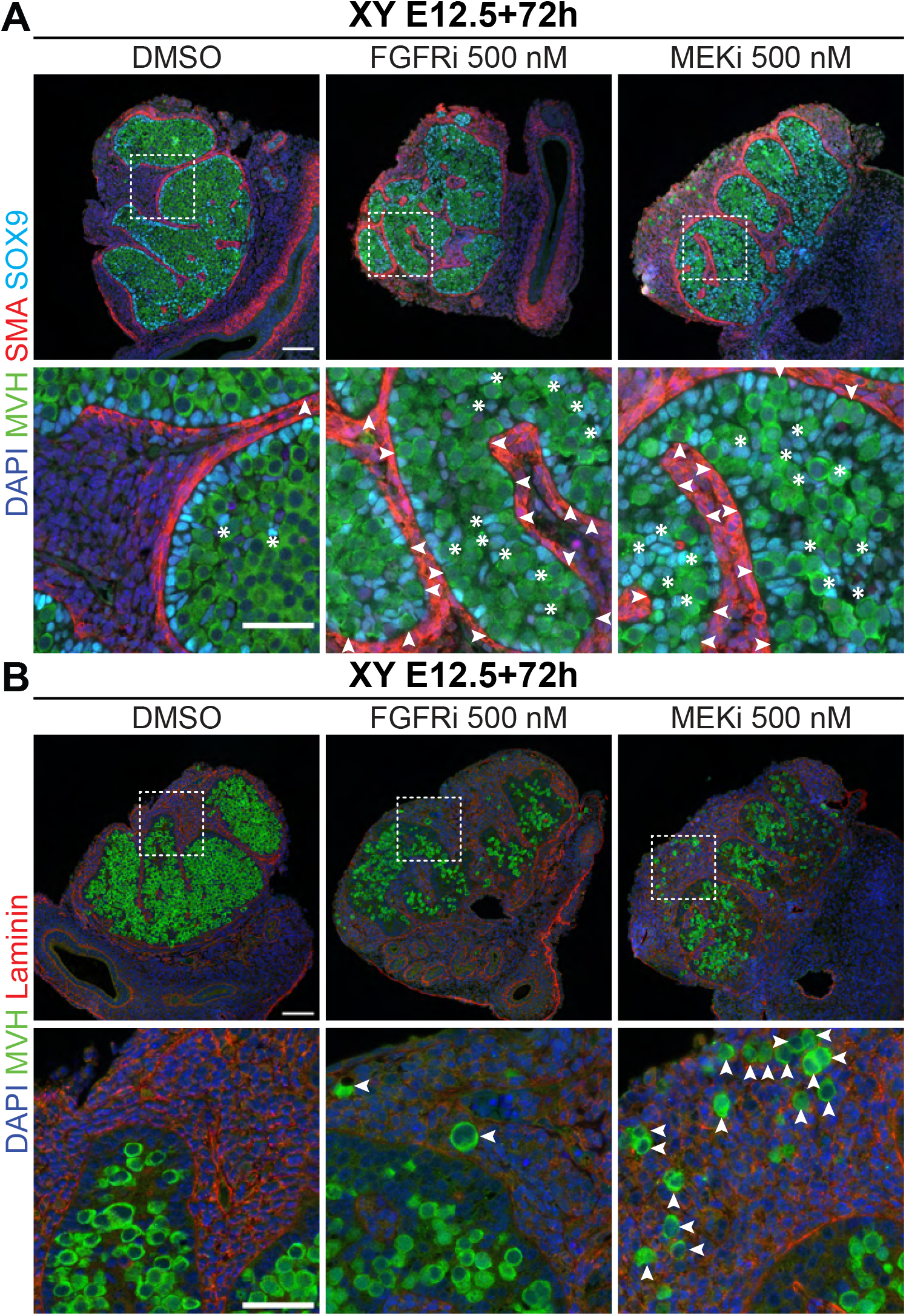
FGF and MEK1/2 signalling is required for normal testis cord formation. Immunofluorescent images of XY E12.5 gonad-mesonephros tissue cultured with DMSO, 500nM FGFRi or 500nM MEKi for 72h showing Sertoli (A) and germ (A-B) cell localisation. DAPI (blue), MVH (green), SMA (red: A) or Laminin (red: B) and SOX9 (cyan: A). Scale bars: top panel 100μm, bottom panel 50μm. Replicates: n=3-4. A: White arrows identify gaps in the Sertoli cell layer at the testis cord basement; white asterisks identify Sertoli cells dispersed within the inner area of the testis cords. B: White arrows identify germ cells localised outside testis cords.

### MEK1/2 signalling is required for male germline differentiation

As MEKi prevented germ cell mitotic arrest, the expression of male germline differentiation markers was assessed using IF and flow cytometry in E12.5 XY gonad-mesonephros samples cultured for 72h with DMSO, 125nM or 500nM FGFRi or MEKi. DPPA4 is expressed in XX and XY germ cells at E12.5, but is upregulated in XY germ cells and repressed in XX germ cells as they differentiate (Gustin et al., 2016b). As expected, DPPA4 was not detected in germ cells of XX gonads but was detected in germ cells of XY E12.5 gonads cultured for 72h with DMSO, and fluorescence appeared more intense in XY E12.5+72h than in E12.5 XY germ cells (Fig 3A, S5A). While DPPA4 germ cell levels were similar in XY E12.5+72h DMSO and FGFRi cultures, DPPA4 intensity appeared lower in MEKi treated samples and comparable to E12.5 XY germ cells (Fig 3A, S5A). Confirming this, flow cytometry revealed that the relative DPPA4 germ cell intensity was 2x higher in XY E12.5+72h DMSO and FGFRi cultures than in E12.5 XY germ cells (*P*<0.0001), but comparable to E12.5 XY germ cells in MEKi-treated gonads (Fig 3B).

**Figure 3:**
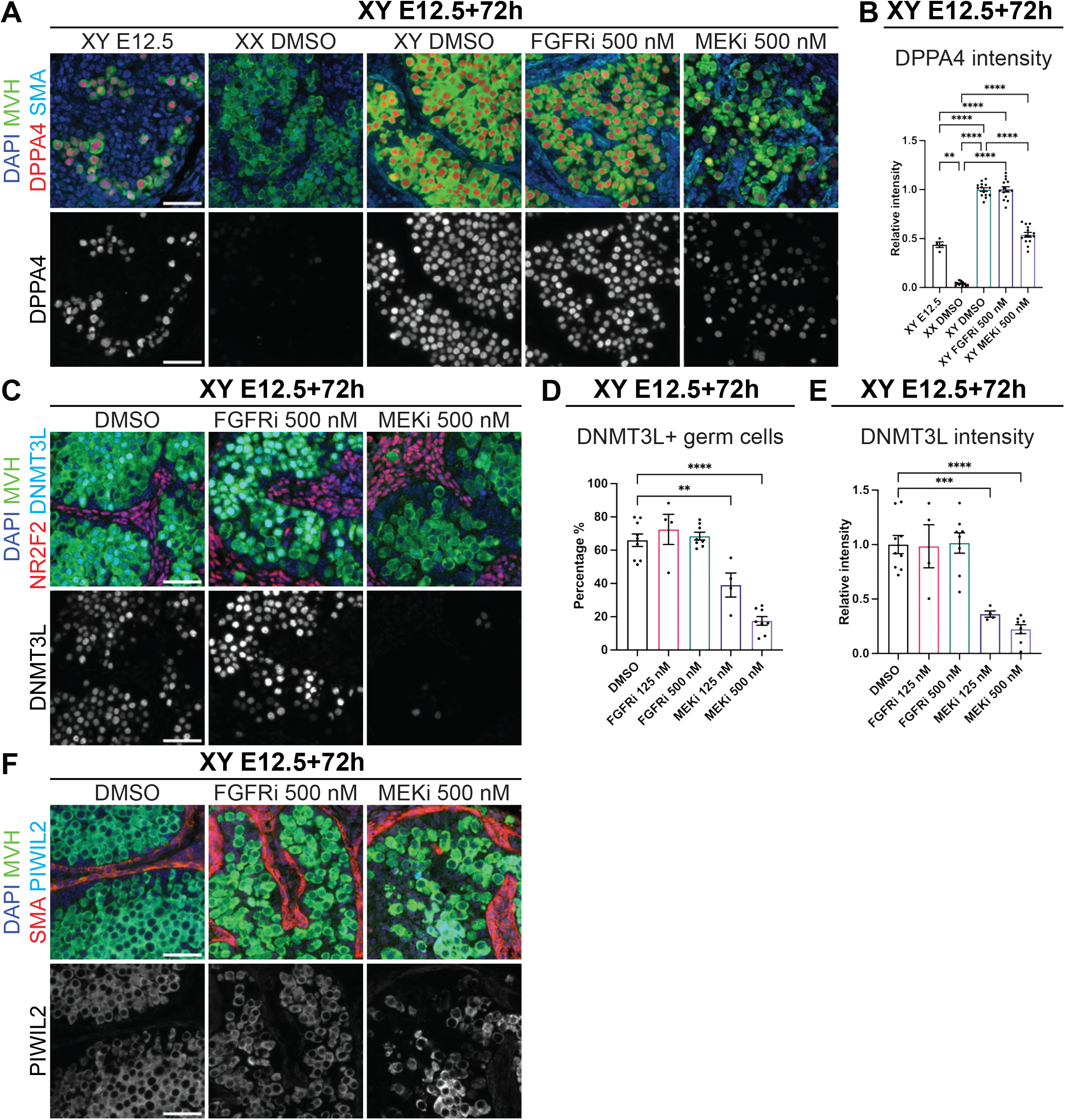
MEK1/2 signalling is required for male germline differentiation. Analysis of E12.5 XY gonad-mesonephros or E12.5 XY or XX gonad-mesonephros cultured for 72h with DMSO, 125 or 500nM FGFRi or MEKi. **A)** Immunofluorescent images demonstrating DPPA4 localisation. Top panel: DAPI (blue), MVH (green), DPPA4 (red), SMA (cyan). Bottom panel: DPPA4 (grey). **B)** DPPA4 staining intensity in germ cells determined by flow cytometry. **C)** Immunofluorescent images demonstrating DNMT3L localisation. Top panel: DAPI (blue), MVH (green), NR2F2 (red), DNMT3L (cyan). Bottom panel: DNMT3L (grey). **D-E)** Percentage DNMT3L positive germ cells (D) and DNMT3L staining intensity (E) determined by flow cytometry. **F)** Immunofluorescent images demonstrating PIWIL2 localisation. Top panel: DAPI (blue), MVH (green), SMA (red), PIWIL2 (cyan). Bottom panel: PIWIL2 (grey). Scale bars: 50μm. Replicates: A, C and F) n=3-4, B) n=4-14, D-E) n=4-9. Statistics: B) Brown-Forsythe and Welch ANOVA with Dunnett’s T3 multiple comparisons, D-E) Ordinary one-way ANOVA with Tukey’s multiple comparison. In B&E; Intensity is relative to E12.5+72h XY DMSO control sample set at 1.0. Error bars: mean ± SEM. Significance between control and treatment: **P<0.01, ***P<0.001, ****P<0.0001.

As the germ cell proliferation and DPPA4 levels indicated that male germ cells failed to properly differentiate in MEKi-treated XY gonads, we examined two additional male germline markers, DNMT3L and PIWIL2. IF and flow cytometry revealed that the majority of germ cells were DNMT3L positive in the DMSO control and FGFRi treated gonads (Fig 3C-E, S5B). In contrast, very few germ cells were DNMT3L positive in MEKi-treated gonads, and the DNMT3L staining intensity was significantly lower than in the DMSO- or FGFRi-treated samples (*P*<0.01, *P*<0.0001; Fig 3C-E, S5B). Similarly, PIWIL2 was expressed at similar levels in germ cells of DMSO- and FGFRi-treated samples but was variable in MEKi-treated samples, with some germ cells staining strongly for PIWIL2 and others negative (Fig 3F, S5C). This was not possible to confirm using flow cytometry because a reliable PIWIL2 flow assay could not be developed.

### MEK1/2 inhibition increased STRA8, but failed to properly induce meiosis in XY germ cells

Since MEKi inhibited mitotic arrest and male germline differentiation, the expression of female germline markers was investigated to determine if FGFRi or MEKi induced female development in XY germ cells. As expected, the pre-meiotic marker STRA8 was detected in the germ cells of XX E12.5+72h DMSO-treated gonads but was not detected in XY DMSO controls (Fig 4A, S6A). While some germ cells appeared very weakly positive for STRA8 in FGFRi treated samples, STRA8 positive germ cells were commonly found in MEKi-treated gonads (Fig 4A, S6A), particularly in germ cells close to the mesonephric-gonadal boundary (Fig S6A). However, while STRA8 staining was localised in the germ cell nucleus in XX controls, it was detected in the germ cell cytoplasm and nucleus in MEKi treatments, indicating that nuclear import-export also regulates STRA8 activity (Fig 4A, S6A). Flow cytometry demonstrated that 74% of germ cells were STRA8 positive in XX DMSO controls, but only 5% and 7% were STRA8 positive in XY DMSO control and FGFRi treated samples (Fig 4B). The proportion of STRA8 positive germ cells in MEKi-treated gonads was 38%, significantly higher than XY controls (*P*<0.0001), but lower than XX controls (*P*<0.0001, Fig 4B).

**Figure 4:**
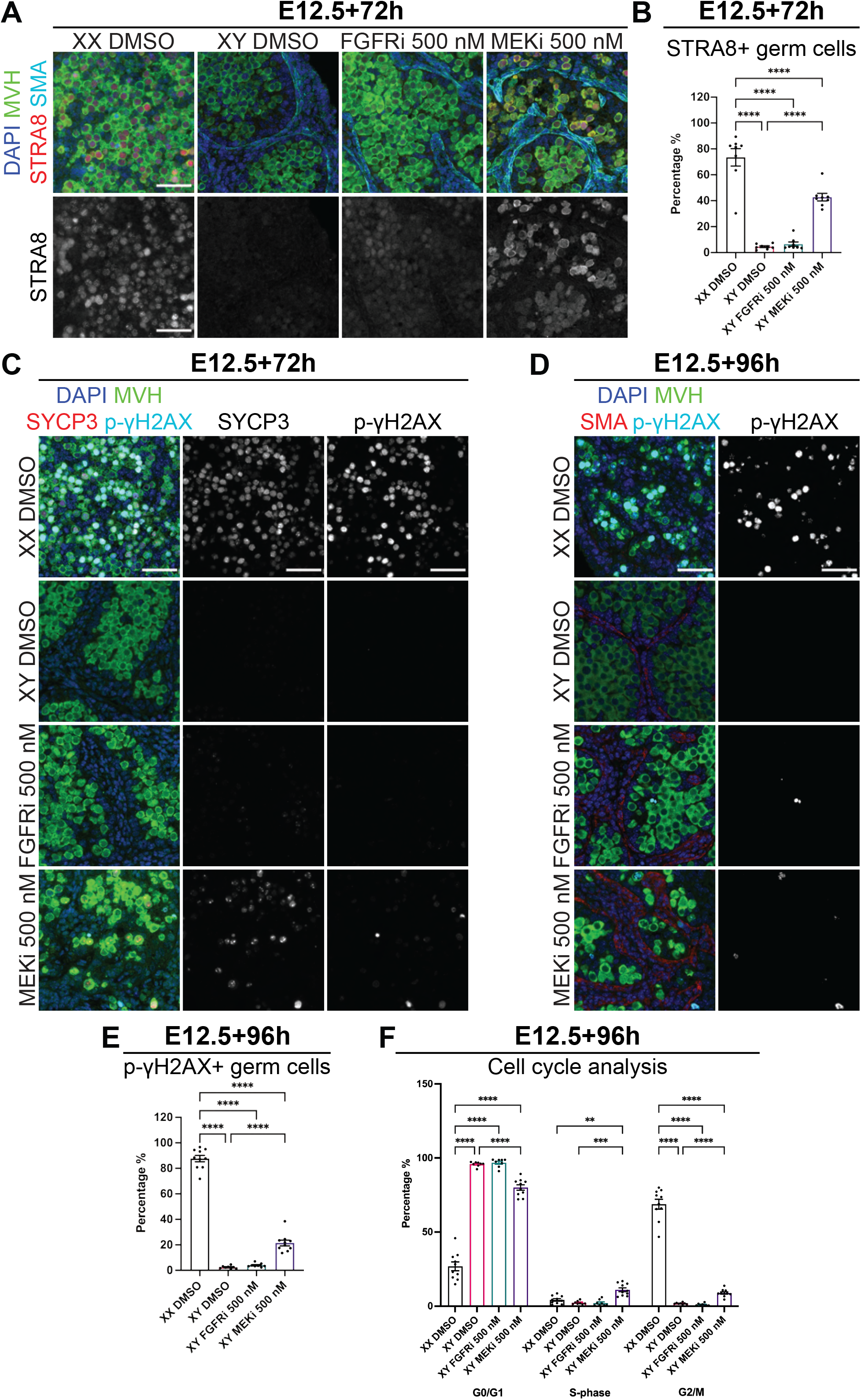
MEK1/2 signalling inhibition permitted *Stra8* expression but failed to effectively induce meiosis in XY germ cells. Analysis of XY or XX E12.5 gonad-mesonephros tissue cultured with DMSO or 500nM FGFRi or MEKi for 72h (A-C) or 96h (D-F). **A)** Immunofluorescent images demonstrating STRA8 localisation. Top panel: DAPI (blue), MVH (green), STRA8 (red), SMA (cyan). Bottom panel: STRA8 (grey). **B)** Percentage STRA8 positive germ cells determined by flow cytometry. **C-D)** Immunofluorescent images demonstrating SYCP3 (C) and phospho-γH2AX (p-γH2AX) localisation. Left panel: DAPI (blue), MVH (green), SYCP3 (red; C) or SMA (red; D) and p-γH2AX (cyan). Middle panel: SCP3 (grey; C). Right panel: p-γH2AX (grey). **E)** Percentage p-γH2AX positive germ cells determined by flow cytometry. **F)** Flow cytometric cell cycle analysis of G0/G1, S-phase and G2/M based on the incorporation of EdU (S-phase) and propidium iodide (DNA content). A, C and D; scale bar: 50μm. Replicates: A, C and D) n=3-4, B) n=8, E-F) n=8-10. Statistics: B) Ordinary one-way ANOVA with Tukey’s multiple comparison, E) Brown-Forsythe and Welch ANOVA with Dunnett’s T3 multiple comparisons, F) Repeated measures two-way ANOVA with Geisser-Greenhouse correction and Tukey’s multiple comparisons. Error bars: mean ± SEM. Significance between controls and treatment: **P<0.05, ***P<0.005, ****P<0.0001.

To determine whether germ cells in FGFRi- or MEKi-treated XY gonads had entered meiosis, gonad sections were triple stained using antibodies specific for SYCP3, phosphorylated γH2AX (p-γH2AX) and MVH (Fig 4C, S6B). SYCP3 and p-γH2AX were detected in most germ cells in 72h XX DMSO controls but not in XY DMSO- or FGFRi-treated gonads (Fig 4C, S6B). A small number of germ cells were positive for SYCP3 in MEKi-treated gonads and a subset also stained for p-γH2AX (Fig 4C, S6B). In addition, rare cells positive for p-γH2AX were detected in XY control, FGFRi and MEKi treated gonads (Fig S6B), however, most did not express MVH and were likely to be apoptotic somatic cells in which p-γH2AX also marks double strand DNA breaks.

To test the possibility that meiotic entry of germ cells in FGFRi- or MEKi-treated gonads was delayed, we cultured E12.5 XX and XY gonad-mesonephros for 96h with DMSO or 500nM FGFRi or MEKi. IF staining revealed that most XX DMSO germ cells were p-γH2AX positive, indicating they had entered meiosis (Fig 4D, S6C). p-γH2AX positive germ cells were rarely detected in XY DMSO- or FGFRi-treated gonads but were more common in MEKi-treated samples (Fig 4D, S6C). Quantification using flow cytometry revealed that 88% of germ cells were p-γH2AX positive in XX DMSO samples while only 2%, 4% and 19% were p-γH2AX positive in XY control, FGFRi and MEKi treated samples, respectively (Fig 4E).

Cell cycle analysis of germ cells from the same gonads using EdU (S-phase) to quantify DNA synthesis and propidium iodide to measure DNA content demonstrated that the majority of germ cells in E12.5+96h cultures were in G2/M in XX DMSO controls, but were in G0/G1 in XY DMSO- and FGFRi treated gonads (Fig 4F). Significantly more germ cells were in G2/M in MEKi than in XY control or FGFRi cultures (*P*<0.0001), but remained less than in XX controls (*P*<0.0001, Fig 4F). Therefore, while MEKi treatment resulted in a significantly greater percentage of p-γH2AX expressing germ cells that were in G2/M, this proportion was substantially lower than in XX controls indicating that meiosis was not properly induced within the normal temporal window following MEK1/2 inhibition.

### The majority of transcriptional divergence occurs after E12.5 in XY and XX germ cells

We next used RNA sequencing to gain greater insight into genome-wide transcriptional changes in FACS isolated *Oct4*-GFP positive germ cells of E12.5 XX and XY gonads (Time 0 controls) and gonads cultured for 24 and 72h with DMSO, FGFRi and MEKi (Fig 5A). Differential expression analysis identified 183 and 234 genes that were expressed higher in XY and XX germ cells at E12.5, respectively (time 0; FDR<0.05; absolute Fold-Change (FC) ≥ 1.5, absolute Log-FC ≥ 0.585; Fig 5B, Table S1.1-1.2). Included in the differentially expressed genes (DEGs) that were higher in E12.5 XY germ cells were a range of Nodal signalling associated genes, including *Nodal, Tdgf1 (Cripto), Lefty1*, *Lefty2, Pitx2* and *Otx2*, which are known to be high in XY germ cells at this time point (Spiller et al., 2012, Miles et al., 2013, Wu et al., 2015, Mayère et al., 2021) (Table S1.1). E12.5 XX germ cells expressed higher levels of BMP target genes, including *Msx1, Msx2, Id1, Id2, Id3, Stra8* and *Gata2,* consistent with observations that BMP2 promotes female germline development (Nagaoka et al., 2020, Mayère et al., 2021) (Table S1.2). However, despite these sex-specific transcriptional differences in E12.5 germ cells, our functional data strongly indicated that these differences were insufficient to ensure male germline commitment as MEK1/2 inhibition at E12.5 substantially disrupted male germline differentiation (Figs 1-3).

**Figure 5:**
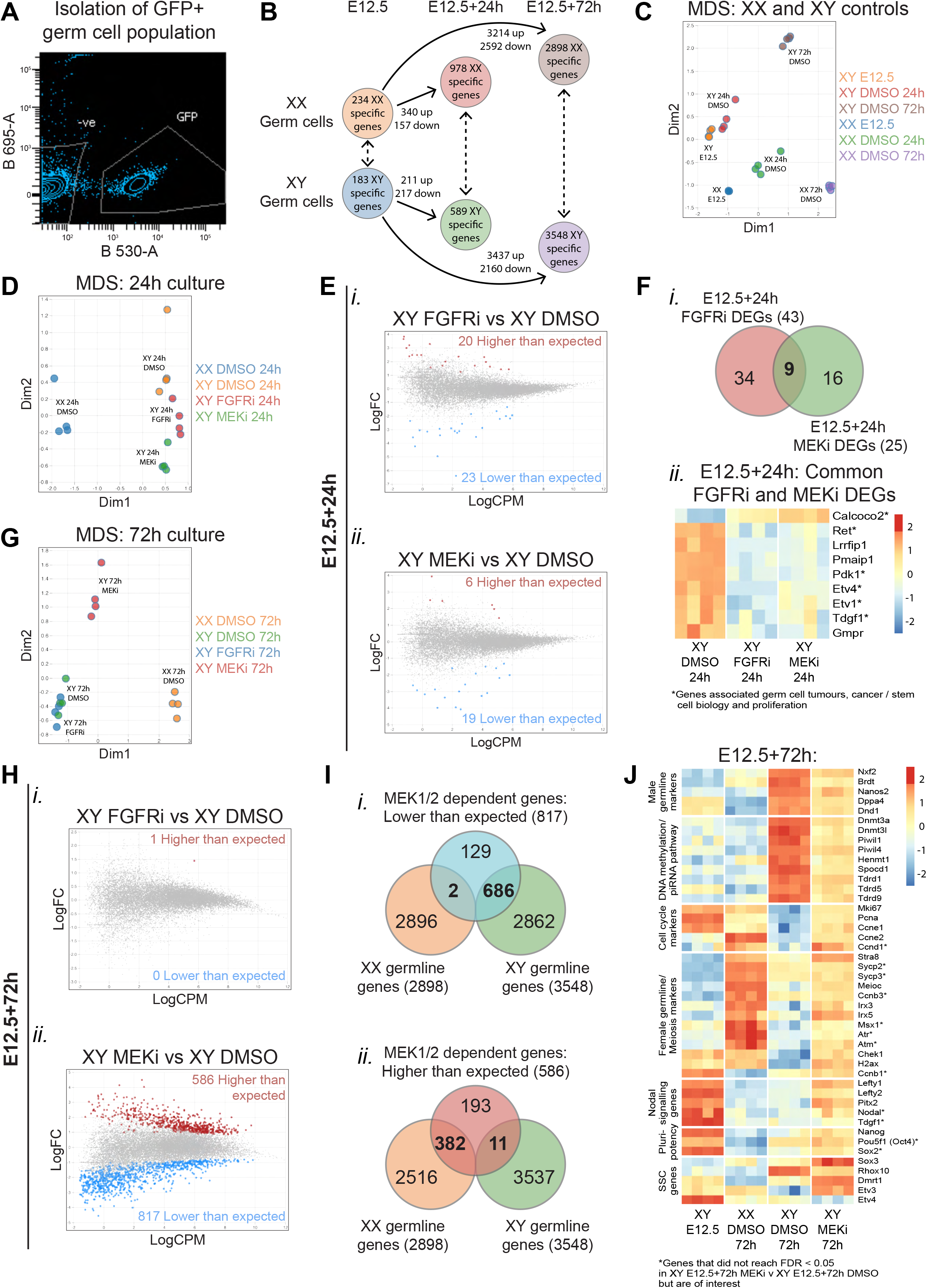
FGF-MEK1/2 signalling supports expression of stem cell associated genes in early germ cells, but only MEK1/2 signalling is required for male germline differentiation. RNA sequencing analysis of germ cells from XX or XY E12.5 gonads, or XX or XY E12.5 gonads cultured for 24 or 72h with DMSO or 500nM FGFRi or MEKi. **A)** Example of FACS scatterplot depicting GFP positive germ cell isolation. **B)** Number of differentially expressed genes (DEGs) between XX or XY E12.5 (time 0) and XX or XY DMSO controls from 24h and 72h cultures. **C)** Multidimensional scaling (MDS) of all control conditions. **D)** MDS of XX and XY gonads cultured for 24h. **E)** Differential gene expression analysis of XY E12.5+24h FGFRi XY (i) or XY E12.5+24h MEKi (ii) vs XY E12.5+24h DMSO. **F)** Venn-diagram of 24h FGFRi and MEKi DEGs (i) and heatmap of common DEGs (ii). Asterisks represent genes associated with germ cell tumours, cancer/stem cell biology and/or proliferation. **G)** MDS of XX and XY gonads cultured for 72h. **H)** Differential gene expression analysis of XY E12.5+72h FGFRi (i) or XY E12.5+72h MEKi (ii) vs XY E12.5+72h DMSO. **I)** Venn-diagram comparing MEKi 72h culture DEGs expressed lower (i) or higher (ii) than expected with XX or XY specific genes identified in B. **J)** Heatmap of DEGs identified in XY E12.5+72h MEKi vs XY E12.5+72h DMSO associated with male germline differentiation, DNA methylation/piRNA pathway, cell cycle, female germline differentiation/meiosis, Nodal signalling, pluripotency and spermatogonial stem cells (SSCs). Asterisks highlight genes associated with cell cycle, meiosis and pluripotency but are not differentially expressed. For all comparisons, genes with FDR <0.05 and |logFC|>0.585 (equivalent to |FC|>1.5) were considered differentially expressed.

To identify male and female transcriptional changes that occurred as a normal part of sex-specific germline differentiation, XX and XY germ cells of DMSO control samples were compared after 24 and 72h of culture. Multidimensional scaling (MDS) revealed that although time 0 XY and XX E12.5 samples were different, they diverged substantially more after 24 and 72h of gonad culture (Fig 5C). Differential gene expression analysis revealed that 211 and 3437 genes were increased, and 217 and 2160 genes were decreased in XY germ cells after 24 and 72h compared to E12.5 (time 0) XY germ cells, respectively (Fig 5B, Table S1.3-1.4). Similarly, 340 and 3214 genes were increased, and 157 and 2592 genes were decreased in XX germ cells compared to E12.5 XX germ cells after 24 and 72h (Fig 5B, Table S1.5-1.6). Together, these data indicated that while male and female germline differentiation progressed in the first 24h, the greatest transcriptional change occurred between 24 (∼E13.5) and 72h (∼E15.5).

To identify genes specifically associated with male and female germline differentiation we compared XY with XX germ cells from DMSO controls. This revealed 589 and 3548 genes higher in XY than XX germ cells after 24 and 72h, (XY germline genes; Fig 5B, Table S1.7-1.8), including male germline genes *Dnmt3l, Dppa4, Nanos2, Piwil1, Piwil2, Piwil4, Tdrd1, Tdrd5* and *Tdrd9*. By comparison, 978 and 2898 genes were higher in XX than XY germ cells after 24 and 72h (XX germline genes; Fig 5B, Table S1.9-1.10), including female germline and meiosis markers *Atr, Atm, Chek1, Ccnb3, H2ax, Irx3, Irx5, Msx1, Id1, Id2, Id3, Sycp2, Sycp3* and *Stra8*, confirming that E12.5 germ cells diverged in the expected sex-specific manner over time.

### FGF and MEK1/2 commonly regulate a subset of genes involved in germ cell tumours, stem cell biology and proliferation

To determine the initial impacts of MEKi and FGFRi on germline development we compared outcomes in XY germ cells cultured with DMSO or 500nM FGFRi or MEKi for 24h. MDS indicated that germ cells from XX E12.5+24h DMSO controls were transcriptionally distinct from all XY groups. XY E12.5+24h DMSO, FGFRi or MEKi cultures were also transcriptionally distinct (Fig 5D) and comparison of XY germ cells from MEKi and FGFRi cultures with DMSO controls revealed 43 and 25 DEGs, respectively (FDR<0.05, |FC|≥1.5; Fig 5E, Table S2.1-2.2). These included 23 and 19 genes that were lower than expected (i.e. for which transcription was not properly activated) and 20 and six genes that were higher than expected (i.e. not properly repressed, or unexpectedly derepressed) in FGFRi or MEKi treatments. Nine genes were commonly dysregulated by MEKi and FGFRi implying that they depend on FGF signalling via MEK1/2 (Fig 5Fi, Table S2.3). Importantly, the direction of change (up- or downregulated) was concordant (roast test *P*=0.00025) for each of the nine common DEGs, with eight expressed lower, and one higher than expected (Fig 5Fii). The simplest interpretation is that the DEGs that were lower than expected were FGF responsive genes that depended on MEK1/2 signalling for their upregulation, and the DEGs that were higher than expected depended on FGF-MEK1/2 for their repression.

Further examination of the genes that were lower than expected revealed six FGFRi-MEKi DEGs associated with stem cell differentiation, cell self-renewal, cancer, cell proliferation and survival and germ cell tumours, including *Etv1* and *Etv4* (Akagi et al., 2015, Gashaw et al., 2005, Oh et al., 2012, Cai et al., 2007), *Pdk1* (Anwar et al., 2021), *Ret* (Miles et al., 2012, Naughton et al., 2006), *Tdgf1* (*Cripto*) (Spiller et al., 2016) and *Calcoco2* (Cui et al., 2021). Few genes commonly associated with sex-specific germline differentiation were represented in the FGFRi and MEKi DEG lists after 24h of culture. However, although unaffected by FGFRi, *Nanos2,* which regulates male germline differentiation (Suzuki and Saga, 2008), was 9.12-fold lower than control in 24h MEKi samples (Table S2.2), consistent with a requirement for MEK1/2 signalling for its upregulation.

### Male specific germline differentiation depends on MEK1/2, but FGF signalling is dispensable

To determine the impacts of FGFRi and MEKi on later stages of male germline differentiation we analysed samples after 72h of culture. Surprisingly, germ cells from 72h XY DMSO and FGFRi cultures were transcriptionally similar (Fig 5G). Although 43 genes were differentially expressed after 24h of FGFR inhibition (FDR<0.05, |FC|≥1.5; Fig 5Ei, Table S2.1), only one DEG was identified after 72h, and was higher than expected (FDR<0.05, |FC|≥1.5; Fig 5Hi, Table S2.4). This may be because *Fgf9* transcription in the testis is normally diminished to very low levels by E14.5 (Bowles et al., 2010). In contrast, MEKi samples were transcriptionally distinct from XX and XY DMSO controls at 72h (Fig 5G), with 817 genes lower and 586 higher than controls (FDR<0.05, |FC|≥1.5; Fig 5Hii, Table S2.5). Of the 817 genes that were lower, 686 were male germline genes (i.e. normally upregulated in the germ cells of XY vs XX E12.5+72h DMSO cultures defined in Fig 5B) and were therefore defined as MEK1/2-dependent male germline genes (Fig 5Ii, Table S2.6). Genes lower than expected included key male germline markers *Dppa4*, *Nanos2*, *Dnd1*, *Nxf2* and *Brdt* (Maldonado-Saldivia et al., 2007, Suzuki and Saga, 2008). Interestingly, the transcriptional levels of *Dppa4* and the germline teratoma gene *Dnd1* (Youngren et al., 2005, Cook et al., 2009) was similar to E12.5 XY germ cells (Fig 5J), consistent with our observations of DPPA4 protein expression in MEKi treated samples (Fig 3A-B). In addition, DNA methylation and piRNA associated genes, including *Dnmt3a*, *Dnmt3l*, *Tdrd1*, *Tdrd5*, *Tdrd9*, *Spocd1*, *Piwil1*, *Piwil4* and *Henmt1* (Sakai et al., 2004, Aravin et al., 2008, Shoji et al., 2009), were not properly upregulated in MEKi treated samples, with levels remaining substantially lower than in XY DMSO-treated controls (Fig 5J). Consistent with persistent germ cell proliferation after MEK1/2 inhibition (Fig 1Eii), genes regulating the G1-S transition, DNA synthesis and germ cell proliferation, including *Mki67, Pcna*, *Ccne1*, *Ccne2* and *Ccnd1*, remained high after 72h of MEKi (Fig 5J).

### MEK1/2 inhibition did not result in overt female germline differentiation

Of the 586 genes that were higher than control in XY germ cells after MEKi (i.e. genes derepressed or not properly repressed via MEK1/2 signalling), 382 were normally expressed at higher levels in differentiating XX than XY germ cells, suggesting feminisation of the germline (Fig 6Iii; Table S2.7). However, of these 382 genes, 218 were expressed at similar levels in E12.5 germ cells (Fig S7), consistent with MEKi blocking germline differentiation rather than increasing feminisation. These included genes marking germ cell proliferation such as *Ccne1, Pcna* and *Mki67* (Miles et al., 2010, Western et al., 2008) (Fig 5J). As MEKi blocked mitotic arrest but most germ cells did not enter meiosis (Fig 4C-F), the simplest explanation is that these genes remained high due to continued germ cell proliferation, rather than female differentiation.

**Figure 6:**
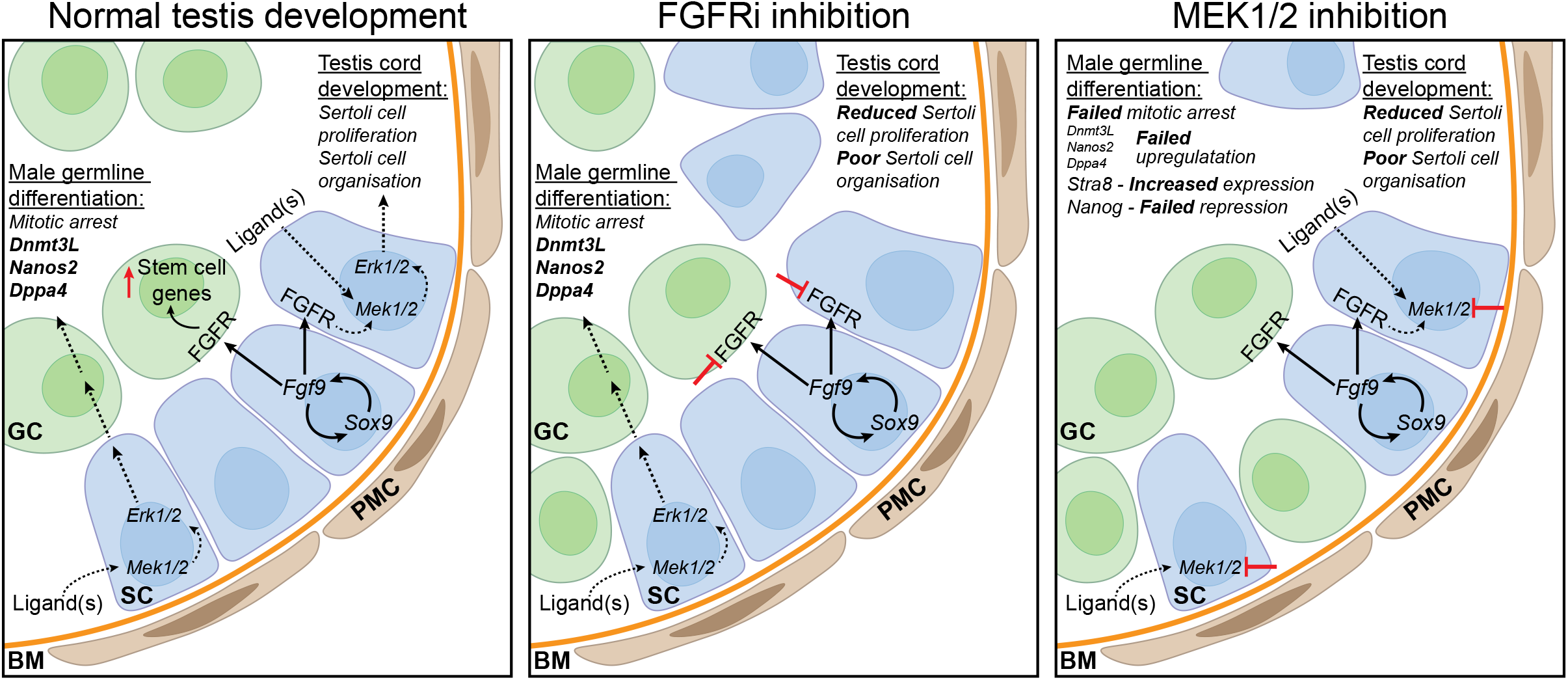
Proposed model for FGF and MEK1/2 signalling in testis and male germline development. In the developing testis, SOX9 and FGF9 promote Sertoli cell proliferation and organisation. It has been proposed that FGF9 drives both germ stem cell characteristics and male germline differentiation. Sertoli cells indirectly promote male germline differentiation, including mitotic arrest and expression of male germline markers such as *Nanos2, Dppa4* and *Dnmt3L.* MEK1/2 signalling inhibition results in failed germ cell mitotic arrest, failed upregulation of male germline markers including *Nanos2,* DPPA4 and DNMT3L and maintained expression of the pluripotency marker, *Nanog.* Although FGF signalling promotes Sertoli cell development and stem cell characteristics in fetal germ cells, it is dispensable for male germline differentiation. We propose that FGF independent MEK1/2 signalling via unknown ligand(s) also promotes Sertoli cell proliferation and organisation to indirectly facilitate male germline development.

MEKi upregulated *Stra8*, *Meioc*, *Irx3* and *Irx5* (Soh et al., 2017, Bowles et al., 2006, Miles et al., 2010, Fu et al., 2018) and some genes that normally increase during female germline development (genes in red box in Fig S7), but the female germline inducing gene *Bmp2* was not upregulated in somatic cells (Fig S3D). Moreover, genes marking meiosis initiation or progression including *Msx1, Sycp2, Sycp3*, *Ccnb3, Atr and Atm* (Miles et al., 2010, Le Bouffant et al., 2011) were not increased by MEKi (Fig 5J). In addition, *Ccnb1,* which is normally repressed in differentiating XX germ cells (Miles et al., 2010), remained high in MEKi cultures (Fig 5J). Other well-known meiosis markers including *Check1* and *H2ax* were transcribed higher in both MEKi and E12.5 XY controls than DMSO controls, so were not informative (Fig 5J). Together, despite higher levels of some female germline markers including *Stra8* and *Meioc*, the low expression of many meiosis markers such as *Atr*, *Atm* and *Msx1* (Fig 5J) was consistent with the limited entry of germ cells into meiosis indicated by analyses of cell cycle and p-γH2AX following MEK1/2 inhibition (Fig 4C-F).

### MEK1/2 inhibition retained germ cells in a relatively undifferentiated state

MEK1/2 inhibition maintained XY germline proliferation and genes including *Dppa4, Dnd1, Ccne1, Ccne2, Ki67, Pcna, H2ax, Chek1* and *Ccnb1* remained at similar levels in E12.5+72h MEKi samples as in E12.5 XY germ cells, indicating that MEKi blocked male germline differentiation (Fig 5J). Consistent with this, the *Nodal* regulatory genes *Lefty1* and *Lefty2* were maintained at higher levels, although *Nodal* and *Tdgf1* were not affected (Fig 5J). While *Nodal* and *Tdgf1* were not affected by MEKi at 72h, the Nodal signalling target *Pitx2* also remained high in MEKi treated samples (Fig 5J).

Key regulators of pluripotency *Sox2* and *Nanog* are expressed in E12.5 XX and XY germ cells but repressed after E13.5 (Western et al., 2010, Maldonado-Saldivia et al., 2007, Bullejos and Koopman, 2004). MEKi maintained *Nanog*, but not *Sox2* transcription in germ cells. *Oct4* remained unchanged, but as *Oct4* transcription is maintained between E12.5 and E15.5 in XY germ cells (Western et al., 2010), this was not informative (Fig 5J). As expected, IF analysis revealed that E13.5 XY germ cells expressed OCT4 and SOX2 protein, but germ cells in XX DMSO+72h cultures were negative (Fig S8). Although OCT4 and SOX2 staining intensity was significantly lower than in E13.5 XY gonads, the proportion of germ cells remaining positive and the staining intensity of these proteins was similar in XY gonads treated with DMSO, FGFRi or MEKi for 72h (Fig S8B-E). In addition, 72h MEKi altered genes associated with germ cell or other tumours (Kanetsky et al., 2011, Taguchi et al., 2021), or SSC function (Raverot et al., 2005, Song et al., 2016) including higher *Dmrt1* and *Sox3* and lower *Rhox10* expression (Fig 5J). Moreover, rather than being down-regulated as normally occurs in differentiating male germ cells, *Etv3* was maintained at levels higher than or similar to E12.5 germ cells, but *Etv4* was repressed (Fig 5J).

### MEK1/2 or FGF signalling inhibition did not cause obvious somatic sex reversal

Our experimental design was to use E12.5 gonad culture to examine germ cells fate within a gonadal environment in the absence of somatic sex-reversal. While collecting germ cells we also isolated the gonadal somatic cells from samples treated with DMSO, FGFRi and MEKi for 24 and 72h. While the bulk of this data will be reported elsewhere, RNAseq analysis revealed no change (based on FDR<0.05, |FC|≥1.5) in the expression of *Foxl2, Rspo1, Bmp2* or *Wnt4,* in samples treated with MEKi or FGFRi compared to DMSO controls (Fig S3D), consistent with IF analysis of SOX9, AMH and FOXL2 (Fig S3A-C). While there was no change in *Cyp26b1* in FGFRi treated samples, MEKi significantly reduced *Cyp26b1* expression (Fig S3D), indicating that this gene may be regulated by MEK1/2, but not directly by FGF signalling. Together, with IF and flow analysis of SOX9, and IF staining of AMH and FOXL2 (Fig S3A-C), these data indicate that FGFRi or MEKi did not result in substantial somatic sex reversal of the gonads.

## Discussion

We have identified a novel and essential role for MEK1/2 signalling in male germ cell differentiation. Our data demonstrate that MEK1/2 signalling is required for XY germ cells to enter mitotic arrest, upregulate a wide range of genes that mark male germline differentiation and repress a range of female germline genes. This occurred in the apparent absence of somatic sex-reversal as SOX9 and AMH expression were normal and ovarian markers were not upregulated. In contrast to MEK1/2, although FGF signalling has been implicated in directly promoting male germline differentiation, our data indicate that FGF signalling is dispensable for germ cell mitotic arrest and male germline differentiation. Instead, our data support a role for FGF-MEK1/2 signalling in regulating genes associated with Nodal signalling and stem cell characteristics in E12.5 XY germ cells, consistent with FGF and Nodal inducing pluripotency in XY germ cells and germ cell tumours, and the well-defined role for FGF in the derivation of pluripotent EG cells (Durcova-Hills et al., 2006, Western, 2009, Bowles et al., 2010, Spiller et al., 2012, Vallier et al., 2009, Mesnard et al., 2006).

MEK1/2 inhibition from E12.5 disrupted germ cell mitotic arrest, maintained expression of genes associated with germ cell proliferation and prevented appropriate upregulation of 686 male germline development genes. A prominent germline signature of MEKi was the deficient upregulation of *de novo* DNA methylation genes and piRNA pathway genes involved in silencing repetitive elements. This implies that the DNA methylation pathway either responds directly to MEK1/2 signalling or is not properly activated due to poor male germ cell differentiation when MEK1/2 is inhibited. Either way, germline DNA methylation is likely to remain low if MEK1/2 signalling is compromised in the germline, potentially allowing de-repression of repetitive elements.

In addition, several genes associated with germline tumours, including members of the Nodal signalling pathway and genes associated with germline tumours in human GWAS and other studies, including *DMRT1* (Turnbull et al., 2010, Taguchi et al., 2021), were expressed in germ cells of MEK1/2 inhibited samples. While we did not observe increased OCT4 or SOX2 protein levels, *Nanog* transcription remained high, germ cell proliferation was sustained, and germline differentiation was inhibited in the absence of MEK1/2 signalling. Combined with low DNA methylation, these factors could render germ cells more susceptible to germline tumours, a possibility that requires further investigation.

A range of studies demonstrate the ability of fetal germ cells to respond to FGF ligands, particularly in isolation from testicular somatic cells (Durcova-Hills et al., 2006, Barrios et al., 2010, Ulu et al., 2017, Sorrenti et al., 2020). Of particular note, FGF signalling (via FGF2, FGF5, FGF9 or FGF10) is required for inducing pluripotency during embryonic germ cell derivation (Durcova-Hills et al., 2006) and FGF9 induces MEK1/2 dependent proliferation in XY germ cells isolated from their somatic counterparts (Ulu et al., 2017). Moreover, FGF and MEK1/2 induce proliferation and underpin an undifferentiated state in SSCs. SSCs express the germline stem cell associated genes *Etv5, Tdgf1, Ret* and, in a more dedifferentiated state, *Nanog* (Ishii et al., 2012, Yang et al., 2021). Consistent with FGF promoting stem cell characteristics in fetal germ cells, we observed a requirement for FGF signalling to regulate 43 genes and MEK1/2 to regulate 25 genes in E12.5 germ cells after 24h of culture. Nine genes were commonly dysregulated by FGFRi and MEKi, suggesting that these genes are regulated by FGF-MEK1/2-ERK1/2 signalling. Six of these genes, including *Etv1, Etv4, Ret, Tdgf1, Pdk1* and *Calcoco,* have been associated with germ cell tumours, stem cells, cell self-renewal, cancer, cell proliferation and cell survival (Akagi et al., 2015, Gashaw et al., 2005, Oh et al., 2012, Cai et al., 2007, Anwar et al., 2021, Miles et al., 2012, Naughton et al., 2006, Spiller et al., 2016, Cui et al., 2021). Although we did not detect pERK1/2 in germ cells using IF, it remains possible that pERK1/2 remained below detection levels and this response occurs directly in germ cells. Interestingly, *Nanos2* is also induced by FGF9 in SSCs and is required to maintain SSCs (Yang et al., 2021, Sada et al., 2012). *Nanos2* was not affected in 24 FGFRi or MEKi cultures but was not upregulated in 72h MEKi cultures. While we did not detect pERK1/2 germ cells, our observations and those of others are consistent with FGF-MEK1/2 signalling priming stemness in germ cells.

Although inhibition of FGFR for 24h from E12.5 resulted in dysregulation of 43 genes, only one gene was dysregulated, and germ cells entered mitotic arrest normally after 72h of FGFRi treatment. In contrast, MEK1/2 inhibition in the same experiment dysregulated 1403 genes, precluded upregulation of the male germline markers DPPA4 and DNMT3L and prevented germ cells from entering mitotic arrest, demonstrating that MEKi disrupted male differentiation in E12.5 XY germ cells. Together, our data indicate that FGF signalling is dispensable, but MEK1/2 signalling is essential for normal differentiation of fetal male germ cells.

It has been suggested that FGF9 directly induces male germline development (Bowles et al., 2010). However, while FGF9 resulted in higher *Nanos2* and *Dnmt3l* transcription in E11.5 XX and XY gonads, experiments in cultured fetal gonads and isolated germ cells have varied (Bowles et al., 2010, Barrios et al., 2010, Gustin et al., 2016b, Ulu et al., 2017). In isolated E12.5 germ cells, 25ng/ml FGF9 either did not significantly increase *Nanos2* or *Dnmt3l* transcription (Bowles et al., 2010) or did increase *Nanos2* (Barrios et al., 2010). A third study found that 25ng/ml FGF9 did not induce *Nanos2* or *Dnmt3l*, but 0.2ng/ml FGF9 did (Ulu et al., 2017). Of note, ectopic FGF9 (20ng/ml) or FGF2 (20ng/ml) promoted proliferation and these factors were able to induce self-renewal genes, including *Nanos2* in SSCs (Ishii et al., 2012, Yang et al., 2021). Moreover, consistent with our observation that FGF9 was required for expression of stem cell related genes in this study, FGF9 increased the pluripotency markers *Oct4* and *Sox2* in E12.5 germ cells (Bowles et al., 2010, Gustin et al., 2016b). Of interest, culture of isolated germ cells or XY gonads with knockout serum replacement, conditions used in some studies, also disrupts germ cell mitotic arrest and transcriptional regulation in developing gonads, enhances germ cell proliferation and favours derivation of pluripotent EGCs (Bowles et al., 2010, Hogg and Western, 2015, Kanatsu-Shinohara et al., 2014, Horii et al., 2003). Together, it appears that while FGF9 is essential in the testis for promoting male somatic cell development and can induce stem cell characteristics in fetal germ cells, the evidence that FGF9 directly promotes male germline development remains limited.

It has also been proposed that low levels of FGF9 (0.2ng/ml) drive male germline differentiation, while high levels (25ng/ml) promote stem cell characteristics (Ulu et al., 2017). We cannot exclude the possibility that residual FGF9 activity drives male germline development in the presence of FGFRi. However, in our study FGFRi completely abrogated FGF9 induced somatic cell proliferation in XX gonads and reduced Sertoli cell proliferation to a similar extent in E12.5 gonads. Moreover, three independent FGFR inhibitors reduced Sertoli cell proliferation to a similar extent but did not affect germ cell mitotic arrest, even when 2500nM FGFRi or 5000nM SU5402, a drug dose which has previously been used to target FGF signalling (Bowles et al., 2010), were used. Furthermore, p38MAPK or PI3K inhibition using two different inhibitors did not affect male germ cell mitotic arrest, indicating that residual FGF9 signalling via p38MAPK or PI3K is unlikely to explain why FGFRi failed to disrupt male germline development.

As FGF9 is at maximal levels at E11.5 in XY gonads and rapidly declines thereafter (Bowles et al., 2010, Jameson et al., 2012b), it remains possible that inhibition of FGFR at E12.5 may have been too late to disrupt male germline development. We were unable to test this possibility as inhibition of FGF signalling at E11.5 is expected to cause somatic sex reversal (DiNapoli et al., 2006, Bowles et al., 2010, Kim et al., 2006, Colvin et al., 2001, Bagheri-Fam et al., 2008, Bagheri-Fam et al., 2017) and would substantially confound the study of sex-specific germ cell development. Nonetheless, MEK1/2 inhibition at E12.5 profoundly disrupted male germline development but FGFRi did not. It is possible that FGF9 acts on germ cells at E11.5 and then MEK1/2 signalling is required at E12.5. However, this would require either a temporal gap between FGFR activation and MEK1/2 signalling, or sustained FGF-MEK1/2 signalling to promote male germline development. Sustained FGF-MEK1/2 signalling would require both FGF and MEK1/2 signalling at E12.5, but our data indicate that only MEK1/2 is required. A temporal gap would require a lag of up to 12-24h (i.e. between E11.5/12 and E12.5) between FGF and MEK1/2-pERK activation. However, FGF induction of MEK1/2 and pERK1/2 typically occurs within 10-15 minutes (Lovicu and McAvoy, 2001, Weyman and Wolfman, 1998), indicating that such a lag is unlikely. Based on past studies and this study, it seems more plausible that FGF9 is dispensable for male germline development in mice, a conclusion that is consistent with normal male germline development in mice lacking *Wnt4* and *Fgf9* (Jameson et al., 2012a).

A model we favour is that FGF9 promotes Sertoli cell and testis development at E11.5 and an additional ligand(s) indirectly promotes male germline fate via MEK1/2 at E12.5 (Fig 6). A combination of signalling by FGF and the other ligand(s) would explain the differing impacts of FGFRi and MEKi on both Sertoli cell proliferation and pERK1/2 in Sertoli cells in this study (Fig 6). While FGFRi completely blocked FGF9-induced proliferation of XX somatic cells, FGFRi had a more modest effect on Sertoli cell proliferation than MEKi. Consistent with this, while MEKi completely abrogated pERK1/2 in Sertoli cells, FGFRi did not and FGFRi reduced Sertoli cell proliferation to a lesser extent than MEKi. As PGD2 is known to facilitate *Sox9* induction and Sertoli cell proliferation, and activate MEK1/2-pERK1/2 (Adams and McLaren, 2002, Wilhelm et al., 2005, Kuroyanagi et al., 2014) it appears likely that FGF9 and PGD2 act together to promote testis development and indirectly drive male germline differentiation.

Deletion of FGF9 causes male to female gonadal sex reversal and germ cells enter meiosis (DiNapoli et al., 2006). While FGF9 treatment of isolated E11.5 germ cells demonstrated that FGF9 can reduce retinoic acid (RA) induced *Stra8* expression, exposure of germ cells to FGF9 in the absence of RA did not significantly affect *Stra8* (Bowles et al., 2010). Moreover, FGFR inhibition in cultured E11.5 XX or XY gonads did not affect *Stra8* expression, but inhibition of CYP26B1 led to robust *Stra8* transcription, presumably due to greater RA availability (Bowles et al., 2010). This effect was enhanced by FGF9 inhibition in XY gonads but not XX gonads (Bowles et al., 2010). While this suggests that FGF9 and CYP26B1 act in concert to inhibit *Stra8* transcription, the effect of FGF9 in this context may also be due to an indirect effect on somatic cells and subsequent *Cyp26b1* transcription in E11.5 gonads rather than a direct effect on germ cells. In our study *Stra8* expression was not affected by FGFR inhibition in E12.5 gonads, perhaps because FGFRi did not reduce *Cyp26b1* transcription and CYP26B1 mediated RA degradation. In contrast, *Cyp26b1* was transcriptionally decreased in MEKi-treated E12.5 testes and *Stra8* transcription and protein were both increased. Despite this, in MEKi-treated samples STRA8 protein was largely localised to the germ cell cytoplasm and *Bmp2* expression was not increased in somatic cells, perhaps explaining why germ cells did not properly enter meiosis. Combined, it seems plausible that the primary role of FGF9 in *Stra8* repression is mediated by ensuring an appropriate testicular environment and consequent *Cyp26b1* expression and *Stra8* repression, rather than a direct effect of FGF9 on germ cells.

Together our study reveals a novel, essential role of MEK1/2 signalling in fetal male germline development, particularly in the promotion of germ cell mitotic arrest, the expression of an appropriate male germline transcriptional program and the activation of *de novo* DNA methylation. While FGF9 may be involved in priming male germline development, it appears to be dispensable for male germ cell differentiation. Moreover, our data suggest that unknown ligand(s) activate MEK1/2 signalling and promote germline differentiation through an indirect mechanism (Fig 6). While PGD2 may be involved, further work is required to determine how this may occur.

## Materials and Methods

### Mouse strains, animal housing, breeding and ethics

Mice were housed at Monash Medical Centre Animal Facility with controlled temperature and humidity, a 12 hours (h) light-dark cycle and food and water available *ad libitum*. Mouse embryos were obtained from inbred 129T2svJ *Oct4*-eGFP males crossed with Swiss females. Females were checked daily for vaginal plugs, with detection of a plug noted as E0.5. Animal work was undertaken in accordance with Monash Medical Centre Animal Facility Animal Ethics Committee approval.

### Organ culture

E12.5 and E13.5 embryos were sexed visually by the presence (male) or absence (female) of testis cords in the gonad. Gonad-mesonephros samples were cultured on 30mm Millicell Biopore membranes with 0.4μm pores (Merck Millipore; PICM03050) in 6-well plates, with each well containing 1400μL media (15mM Hepes, 0.1mM non-essential amino acids, 1mg/mL N-acetylcysteine, 1X penicillin/streptomycin, 55μM beta-mercaptoethanol and 10% fetal calf serum in DMEM/F12 with Glutamax). PBS was placed in between the wells to maintain humidity. Gonads were cultured in media containing DMSO (vehicle control, was used at a dilution of ≥1/5,000, as appropriate for the concentration of each drug dilution), 125, 250, 500 1000nM of BGJ398 (FGFRi; SelleckChem, HY-13241), PD0325901 (MEKi; SelleckChem, S1036), GSK1059615 (PI3Ki; MedChem Express, HY-12036), PH797804 (p38i; MedChem Express, HY-10403). Additional inhibitors for FGFR (AZD4547; SelleckChem, S2801 and SU5402; MedChem Express, HY-10407), p38 (Ralimetinib dimesylate; MedChem Express, HY-13241) and PI3K (PF-04691502; MedChem Express, HY-15177) signalling were also used to ensure consistency of outcomes with each of the primary inhibitors used. All inhibitors were selected based on their high-specificity, potency and advancement in clinical trials (Table 1). Gonad-mesonephric complexes were randomly allocated to each culture treatment condition and cultured for 24, 48, 72 or 96h in 5% CO_2_ at 37°C, with media refreshed daily. To facilitate analysis of cell proliferation, 5-ethynyl-2’-deoxyuridine (EdU) was added to each sample for the final two hours of culture at a final concentration of 20μM. After culture, gonads were photographed under brightfield and fluorescence optics, then processed for flow cytometry, IF or fluorescent activated cell sorting (FACS). For gonad only flow cytometric experiments and all experiments involving FACS purification of germ and somatic cells, gonads were dissected away from mesonephros at the end of the culture period.

### Flow cytometry

Gonad collection, dissociation, fixation, staining and flow cytometry were performed essentially as described previously (Wakeling et al., 2013), using eGFP or antibodies specific for MVH, DPPA4, SOX9, DNMT3L, STRA8 or H2AX. Mesonephros or limb samples were used as germ cell negative controls to set gates for eGFP, MVH or DPPA4 and E12.5 female gonads were used as a negative control to set gates for SOX9 and DNMT3L. Cultured male gonads were used as a negative control to set gates for STRA8 and H2AX. Representative gating and negative control gates can be found in S1. A rabbit IgG antibody was used as a negative control for determining staining intensities using specific rabbit antibodies. Primary antibodies used are listed in Table 2. Secondary antibodies used include Alexa Fluor Donkey anti Goat 488 (Thermo-Fisher, A11055), Alexa Fluor Donkey anti Goat 647 (Thermo-Fisher, A31573), Biotin Donkey anti Rabbit (Thermo-Fisher, A16027) and Biotin Donkey anti Goat (Thermo-Fisher, A16009). Cell cycle analysis was performed as previously described (Wakeling et al., 2013), with germ cells or Sertoli cells identified by their expression of MVH or SOX9, respectively. Cells were stained with 20μg/mL of propidium iodide, enabling quantitation of cellular DNA content. Proliferation was measured by gating EdU positive cells to identify cells in S-phase, while cells in G0/G1 or G2/M were respectively identified by DNA contents estimated as 2n or 4n in the EdU negative population compared to DMSO controls. All flow cytometry was performed on a BD FACS Canto II analyser (BD, Biosciences) and data analysed with FlowJo (v10.7.2) and Graphpad Prism (v9.2.0). Relative antibody staining intensities were calculated by removing background staining in negative control samples and normalising levels to the XY DMSO control set at 1.0. Data represents 3-16 biological replicates and statistical significance was determined using one-way ANOVA with Tukey’s multiple comparisons or unpaired two-tailed t-test, where appropriate. If variance was unequal, a non-parametric Brown-Forsythe and Welch ANOVA with Dunnett’s T3 multiple comparisons was used. *P* values <0.05 were considered significant.

**Table 2:** Antibodies for flow cytometry (F) and immunofluorescence (IF)

| Protein | Source and catalogue # | Species | Dilution |
| --- | --- | --- | --- |
| Mouse Vasa Homologue (MVH) | R&D Systems, AF2030 | Goat | F: 1/100<br>IF: 1/400 |
| DPPA4 | R&D Systems, AF3730 | Goat | F: 1/100<br>IF: 1/400 |
| OCT4 | Santa Cruz, sc8628 | Goat | IF: 1/400 |
| AMH | Santa Cruz, sc68886 | Goat | IF: 1/200 |
| MVH | Cell Signalling Technology, 8761S | Rabbit | IF: 1/400 |
| SOX9 | Sigma-Aldrich, AB5535 | Rabbit | F: 1/200<br>IF: 1/1000 |
| DNMT3L | Cell Signalling Technology, 13451S | Rabbit | F: 1/100<br>IF: 1/100 |
| STRA8 | Abcam, ab49405 | Rabbit | F: 1/500<br>IF: 1/400 |
| Phospho-γH2AX | Cell Signalling Technology, 9718S | Rabbit | F: 1/100<br>IF: 1/800 |
| IgG | Cell Signalling Technology, 3900S | Rabbit | Equal to primary antibody used |
| Laminin | R&D Systems, L9393 | Rabbit | IF: 1/200 |
| PIWIL2 | Cell Signalling Technology, 5940S | Rabbit | IF: 1/200 |
| Phospho-ERK1/2 | Cell Signalling Technology, 4370S | Rabbit | IF: 1/200 |
| FOXL2 | A gift from A/Prof Dagmar Wilhelm | Rabbit | IF: 1/500 |
| SYCP3 | Abcam, ab97672 | Mouse | IF: 1/200 |
| Smooth Muscle Actin (SMA) | Sigma-Aldrich, A2547 | Mouse | IF: 1/1000 |
| NR2F2 | R&D Systems, PP-H7147-00 | Mouse | IF: 1/400 |
| SOX2 | Cell Signalling Technologies, 4900S | Mouse | IF: 1/100 |

### Tissue fixation, embedding, immunofluorescence and image analysis

Gonads were fixed in 4% paraformaldehyde (PFA) in PBS overnight at 4°C. Samples were washed three times in PBS before 70% ethanol processing, and embedded in paraffin. 4μm sections were cut in a compound series (typically four slides prepared per gonad sample with around 6 sections collected/slide), mounted on Superfrost Plus slides and dried at least overnight before antibody incubation. Antigen retrieval was conducted using Dako Citrate buffer (pH 6.0) for 30 minutes at 98°C in a PT Link rinse station (Dako). Tissue sections were blocked in PBTx containing 5% BSA (Merck, A9647) and 10% donkey serum (Sigma; D9663) for 1 hour at RT. Primary antibody (Table 2) diluted in PBTx containing 1% BSA was left to incubate overnight at 4°C or for 2h room temperature (RT). Slides were incubated for 1h at RT in the dark in secondary antibody (Alexa Fluor, Thermo-Fisher, Donkey anti Goat 488 A11055 or Donkey anti Rabbit 488 A21206; Donkey anti Rabbit 647 A31573 or Donkey anti Mouse 647 A31571; Donkey anti Mouse 594 A21203; Donkey anti Goat 594 A11058 or Donkey anti Mouse 555 A31570) diluted at 1/300 in PBTx containing 1% BSA. Slides were mounted in ProLong Gold containing DAPI (Thermo-Fisher, P36931). Confocal images were taken using a Nikon C1 Confocal microscope, with images taken using either 10x lens or 40x oil immersion lens, or slides were scanned using a VS120 Virtual Slide microscope (Olympus), collecting single optical sections of the whole area for each section on the slide. Image analysis was conducted with QuPath (v0.3.0) (Bankhead et al., 2017). Data from at least three representative tissue sections from four biological replicates were averaged. Relative antibody staining intensity was calculated by normalising levels to the XY DMSO control set at 1.0. Data represents four biological replicates and statistical significance was determined with Graphpad Prism (v9.2.0) using one-way ANOVA with Tukey’s multiple comparisons. If variance was unequal, a non-parametric Brown-Forsythe and Welch ANOVA with Dunnett’s T3 multiple comparisons was used. *P* values <0.05 were considered significant.

### Fluorescent activated cell sorting (FACS) of germ and somatic cells

After organ culture, the mesonephros was dissected from the gonads. Six to 15 gonads were pooled for each sample and were dissociated in trypsin. Germ and somatic cell populations were isolated as previously described (Western et al., 2008, Wakeling et al., 2013) using the BD FACSAria^TM^ Fusion cell sorter. GFP positive germ and GFP negative somatic cell populations were isolated from E12.5 XX and XY control gonads cultured for 24 or 72h in DMSO or XY E12.5 gonads cultured for 24 or 72h in DMSO, FGFRi or MEKi (n≥4 for each treatment/group). Germ cells were defined as GFP positive, with dead propidium iodide positive germ cells excluded.

### RNA-sequencing library construction and sequencing

RNA was isolated from 3-45 x 10^4^ FACS-sorted germ cells using Macherey-Nagel NucleoSpin® RNA XS extraction kit (Scientifix, 740902.50) following manufacturer instructions. RNA quantity and RNA integrity (RIN) were assessed using Qubit and Bioanalyzer (Agilent Technologies). Libraries were prepared with 30ng of RNA from germ cells with RIN values greater than 7. The library was constructed as previously described by the MHTP Medical Genomics Facility (Grubman et al., 2021). Briefly, during initial poly(A) tail priming, an 8bp sample index along with a 10bp unique molecular identifier (UMI) was added. Samples were pooled and amplified with a template switching oligonucleotide. The Illumina P5 and P7 were added by PCR and Nextera transposase, respectively. The forward read (R1) utilises a custom primer to sequence into the index while the reverse read (R2) uses a standard R2 primer to sequence the cDNA in the sense direction. I7 indexes were added to enable parsing of sample sets. Sequencing was performed on NextSeq2000 (Illumina) following Illumina protocol 1000000109376 v3.

### Data pre-processing

FASTQ files were demultiplexed and mapped using scPipe (Tian et al., 2018) and Rsubread (Liao et al., 2019) (Bioconductor) packages in R studio. Briefly, FASTQ files were reformatted with *sc_trim_barcode* to incorporate barcode information from read 1 into read 2 into the header. Reads were aligned to a reference mouse genome (GENECODE GRCm39) using Rsubread. Reads were assigned to annotated exons with sc*_exon_mapping*, data were demultiplexed using *sc_demultiplex* and a gene count matrix was generated with UMI deduplication using *sc_gene_counting*. Gene count matrices from each set were combined into a DGEList object for analysis.

### Downstream analysis of RNA-sequencing data

Differential gene expression was assessed using the Limma (Ritchie et al., 2015), Glimma (Su et al., 2017) and edgeR (Robinson et al., 2009) Bioconductor packages following a previously established workflow (Law et al., 2016). Briefly, gene count data was loaded into R studio and genes were annotated with any duplicates removed. Raw counts were transformed into counts per million (CPM). Lowly expressed genes were removed using the *filterByExpr* function in edgeR, and gene expression distributions were normalised using trimmed mean of M-values (TMM) method (Robinson and Oshlack, 2010). Multidimensional scaling (MDS) plots were generated to visualise sample clustering. Since there were many samples present, groups were subdivided into smaller groups to visualise clustering between different culture treatments and culture periods more clearly. Heteroscedasticity was removed from the count data with *voomWithQualityWeights* (Liu et al., 2015). Linear modelling and empirical Bayes moderations was used to test for differential expression. As there were many differentially expressed genes (DEGs) identified following empirical Bayes moderations, to identify DEGs of biological relevance, a log_2_fold-change cut-off was set at greater/less than 0.585 (equivalent to a FC of 1.5) using *treat* (McCarthy and Smyth, 2009). Genes were considered differentially expressed if they met the logFC cut-off and had a false discovery rate (FDR) adjusted p-value less than 0.05. DEGs identified in XY E12.5+24h FGFRi vs XY E12.5+24h DMSO (24h FGFRi DEGs) and XY E12.5+24h MEKi vs XY E12.5+24h DMSO (24h MEKi DEGs) were compared, with genes commonly affected by FGFRi and MEKi assessed using Roast (Wu et al., 2010). Heatmaps were generated using ClustVis (Metsalu and Vilo, 2015).

## Supporting information

Blucher etal Supplementary Table 1

Blucher etal Supplementary Table 2

## Declarations

### Funding

This work was supported by grants and research funding from:

Australian Research Council Discovery Project DP210102342 (PSW, MER)

Hudson Institute of Medical Research (PSW)

Victorian Government’s Operational Infrastructure Support Program

Monash University Postgraduate Student Award to ROB

### Author contributions

Conceived and designed experiments: ROB, PSW

Performed experiments and/or analysed data: ROB, EGJ, PSW

Bioinformatic analyses: ROB, RSL

Writing - original draft: ROB, PSW

Writing - review & editing: ROB, PSW, MER

All authors critically read and approved the manuscript.

### Ethics approval and consent to participate

All animal work was undertaken in accordance with Monash University Animal Ethics Committee (AEC) approvals.

### Data and materials availability

With exception of the RNA sequencing data generated in this study, all data are available in the main text and the Supplementary Materials. The RNA sequencing data have been deposited in the Gene Expression Omnibus (GEO) and are publicly available via accession number GSE221453.

## Acknowledgments

We thank Associate Professor Dagmar Wilhelm and Associate Professor Craig Smith for critical comments on the manuscript, Monash Animal Research Platform staff for assistance with mouse care, Monash Histology Platform for assistance with slide scanning, the Monash Micro Imaging Facility and MHTP Medical Genomics Facilities for assistance and technical advice and Peter Hickey for advice regarding bioinformatic analysis.

## Supplementary Tables and Figures

**Tables S1 and S2**

Supplementary Tables 1 and 2 contain gene lists generated from RNA sequencing analyses performed in this study.

**Figure S1:**
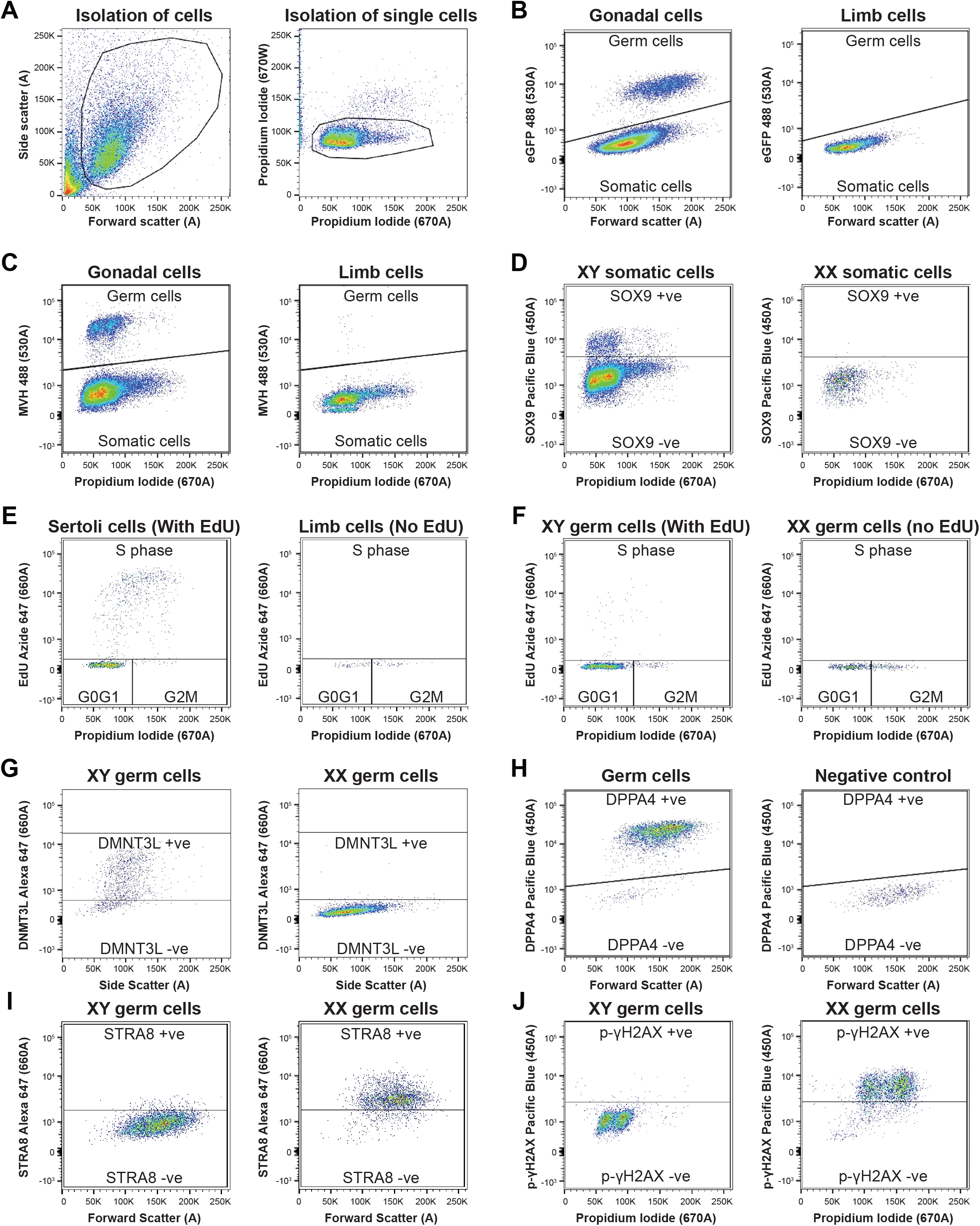
Representative plots depicting gating for antibodies used in flow cytometric analysis. A = Area, W = Width. **A)** Gates used to separate cells from debris (left) and to isolate single cells based on propidium iodide (PI) staining (right). **B)** Germ and somatic cells populations were identified by detecting Oct4GFP transgene (left). E12.5 mouse limb or mesonephros cells were used as a negative control (right). **C)** Germ and somatic cells populations were identified by detecting MVH staining (left). E12.5 mouse limb or mesonephros were used as a negative control (right). **D)** Sertoli cells were identified based on SOX9 staining (left). XX somatic cells were used as a negative control (right). **E-F)** Incorporation of EdU was used to identify proliferating Sertoli cells (E) or germ cells (F), with PI incorporation used to determine individual cell DNA content (left). E12.5 limb or mesonephros (E) or E12.5 XX germ cells not exposed to EdU (F) were used as a negative control for EdU (right). **G)** DNMT3L+ germ cells identified with DNMT3L staining (left). XX germ cells were used as a negative control (right). **H)** DPPA4+ germ cells were identified with DPPA4 staining (left). Cells not stained for DPPA4 were used as a negative control (right). **I-J)** E12.5+72h XY DMSO germ cells were used as a negative control for STRA8 (I) or p-γH2AX (J) staining (left). E12.5+72h XX DMSO germ cells were used as a positive control (right).

**Figure S2:**
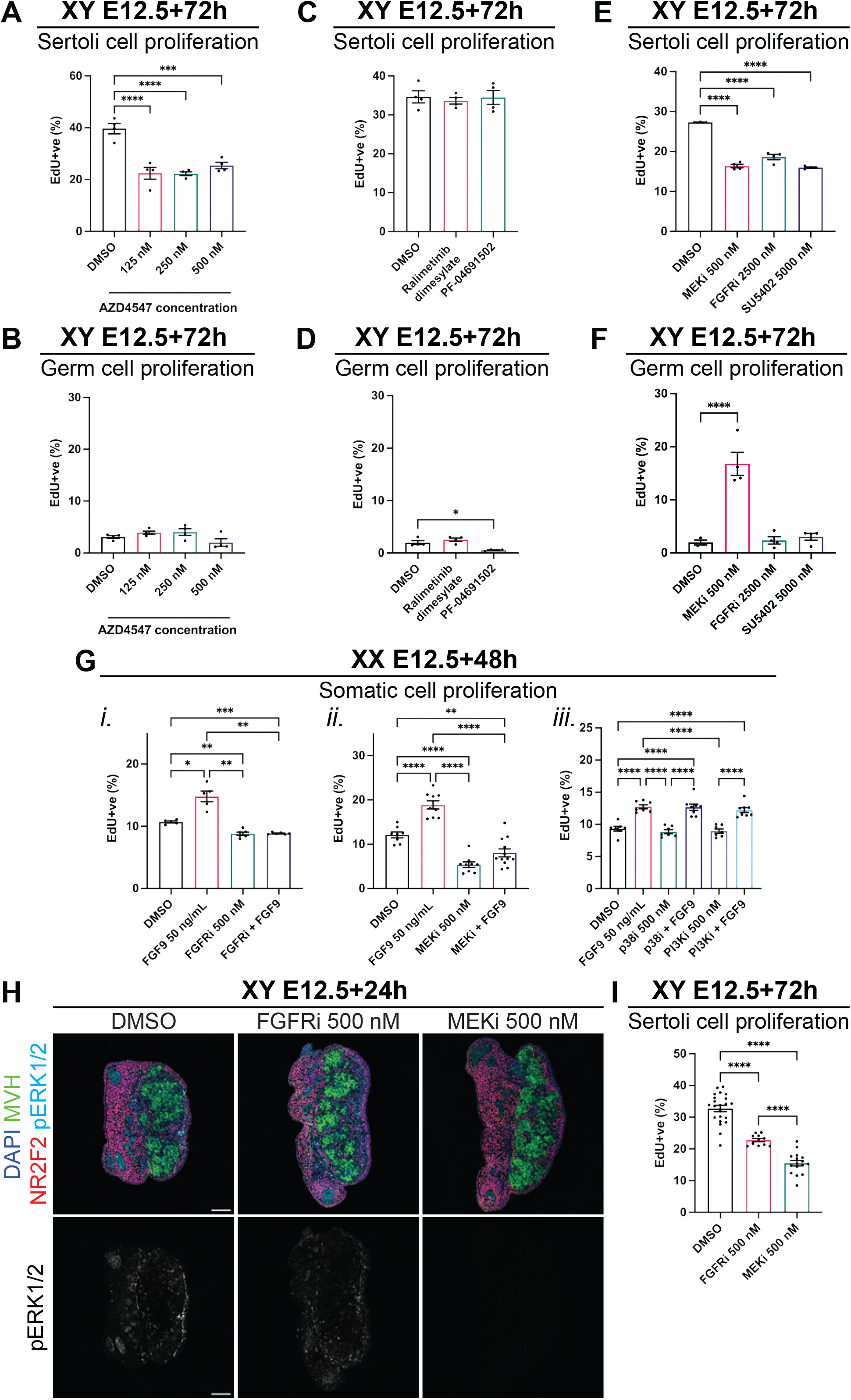
FGFR or MEK1/2 inhibition reduced Sertoli cell proliferation, but only MEK1/2 inhibition disrupted germ cell mitotic arrest. **A-F)** Flow cytometric analysis of Sertoli (A, C and E) or germ (B, D and F) cell proliferation based on EdU incorporation in XY E12.5 gonad-mesonephros tissue cultured for 72h with DMSO, 125, 250 or 500nM of FGFR inhibitor, AZD4547 (A-B), 500nM of p38 inhibitor, ralimetinib dimesylate or 500nM of PI3K inhibitor, PF-04691502 (C-D) or 500nM of MEKi, 2500nM of FGFRi or 5000nM of FGFR inhibitor, SU5402 (E-F). **G)** Flow cytometric analysis of gonadal somatic cell proliferation identified by EdU incorporation in E12.5 XX gonads/mesonephros tissue cultured for 48h with DMSO, FGF9 (50ng/mL), 500nM of FGFRi (i), MEKi (ii), p38i or PI3Ki (iii) and FGF9 + FGFRi (i), FGF9 + MEKi (ii), FGF9 + p38i or PI3Ki (iii). **H)** Wide view immunofluorescent images of E12.5 gonad-mesonephros tissue cultured for 24h with DMSO, 500nM of FGFRi or MEKi demonstrating MEK1/2 signalling activity. Top panel: DAPI (blue), MVH (green), NR2F2 (red), pERK1/2 (cyan). Bottom panel: pERK1/2 (grey). Scalebar represents 100μm. **I)** Flow cytometric analysis of Sertoli cell proliferation based on EdU incorporation in XY E12.5 gonad-mesonephros tissue cultured for 72h with DMSO, 500nM FGFRi or MEKi. Replicates: A-D) n=4, E-F) n=3-4, Gi) n=5-6, Gii) n=8-12, Giii) n=11-21. Statistics: A-F,Gii-Giii) Ordinary one-way ANOVA with Tukey’s multiple comparison, Gi and I) Brown-Forsythe and Welch ANOVA with Dunnett’s T3 multiple comparisons. Error bars: Mean ± SEM. Significance between controls and treatment: *P<0.05, **P<0.01, ***P<0.001, ****P<0.0001.

**Figure S3:**
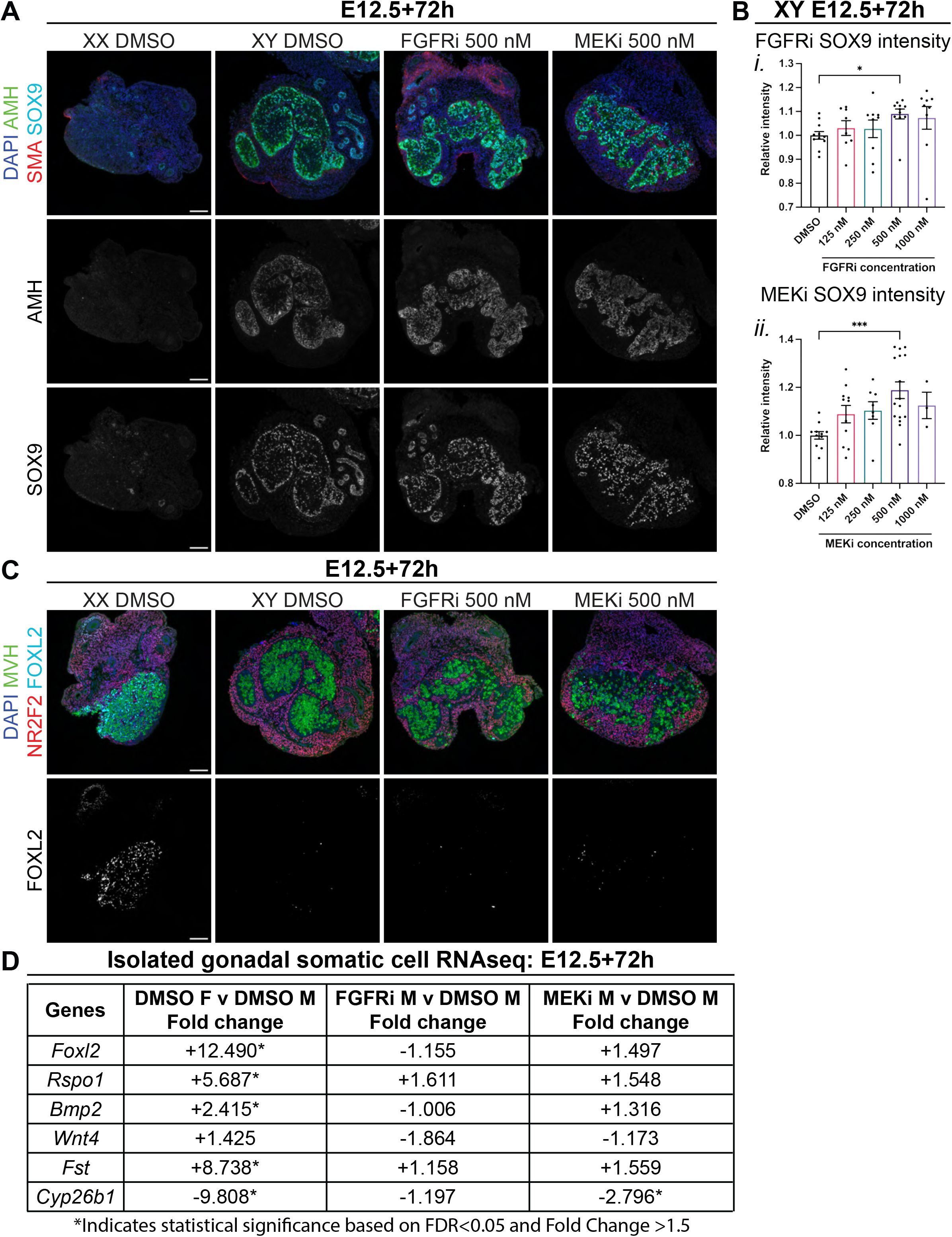
FGF and MEK1/2 inhibition from E12.5 does not cause sex reversal of the gonads. Analysis of XY E12.5 or XX E12.5 gonad-mesonephros tissue cultured with DMSO or 500nM of FGFRi or MEKi for 72h. **A)** Immunofluorescent images demonstrating AMH and SOX9 staining. Top panel: DAPI (blue), AMH (green), SMA (red), SOX9 (cyan). Middle panel: AMH (grey). Bottom panel: SOX9 (grey). **B)** SOX9 staining intensity in Sertoli cells determined by flow cytometry. **C)** Immunofluorescent images demonstrating FOXL2 staining. Top panel: DAPI (blue), MVH (green), NR2F2 (red), FOXL2 (cyan). Bottom panel: FOXL2 (grey). **D)** RNA sequencing results in isolated gonadal somatic cells following 72h culture of key female and male gonadal somatic cell markers including *Foxl2*, *Rspo1*, *Bmp2*, *Wnt4*, *Fst* and *Cyp26b1.* Data shows the fold change between XX E12.5 DMSO v XY E12.5 DMSO, XY E12.5 FGFRi v XY E12.5 DMSO and XY E12.5 MEKi v XY E12.5 DMSO. Plus (+) or minus (–) symbol indicates increased or decreased expression, respectively. Asterisks indicates statistical significance based on FDR<0.05 and FC >1.5. Scale bar represents 100μm. Replicates: A and C) n=3-4, Bi) n=8-11, Bii) 3-16. Statistics: Bi) Welch ANOVA with Dunnett’s T3 multiple comparisons, Bii) Ordinary one-way ANOVA with Tukey’s multiple comparison. In B; Intensity is relative to DMSO control sample set at 1.0. Error bars: Mean ± SEM. Significance between controls and treatment: *P<0.05, ***P<0.001.

**Figure S4:**
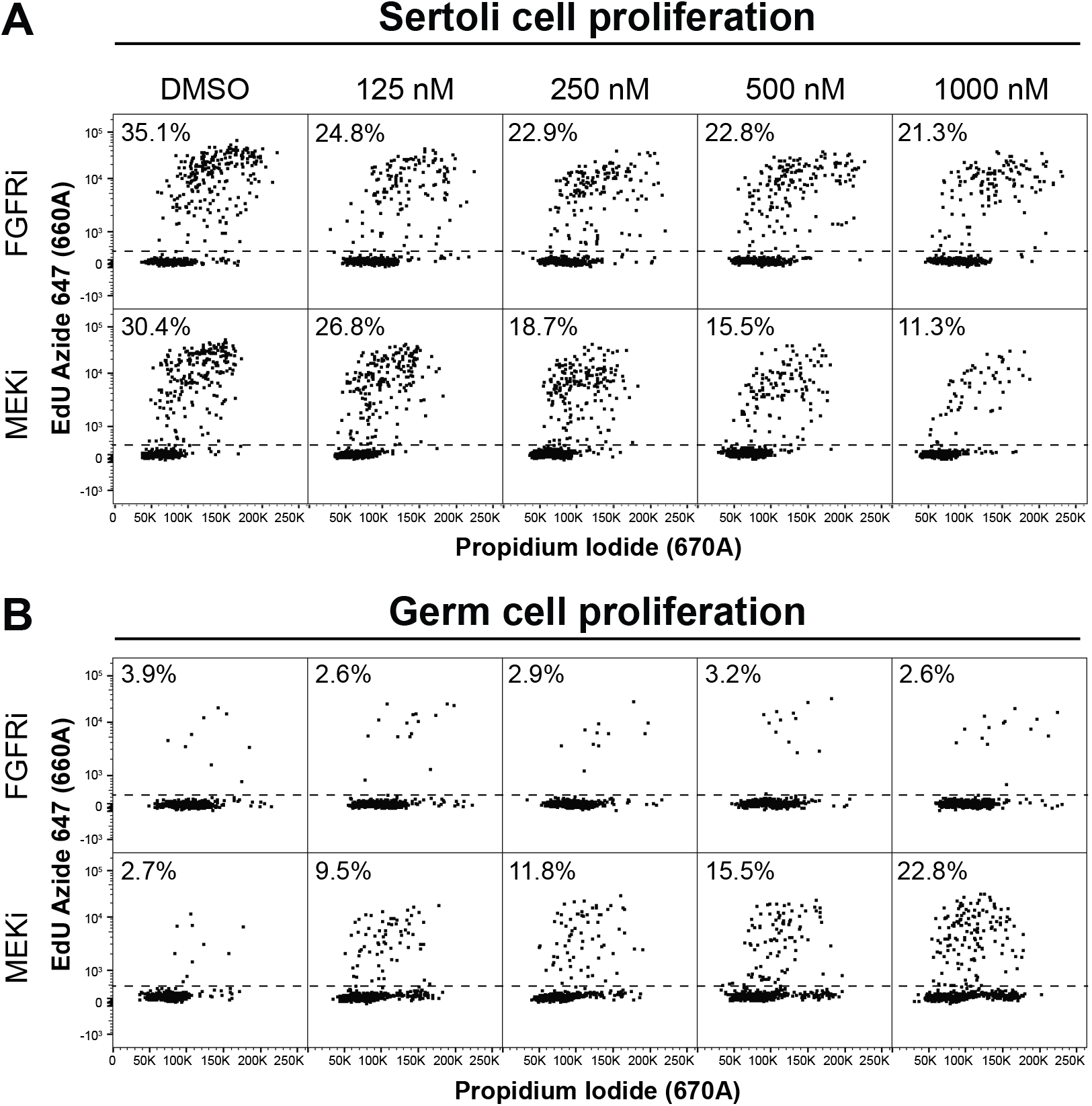
Flow cytometric scatterplot depicting Sertoli and germ cell proliferation. Flow cytometric scatterplots of XY E12.5 gonad-mesonephros tissue cultured in DMSO, 125, 250, 500 or 1000nM of FGFRi or MEKi for 72h showing the percentage EdU incorporation in Sertoli (A) or germ (B) cells. Percentage in top left corner of each graph represents the average proportion of Sertoli (A) or germ (B) cells in each treatment.

**Figure S5:**
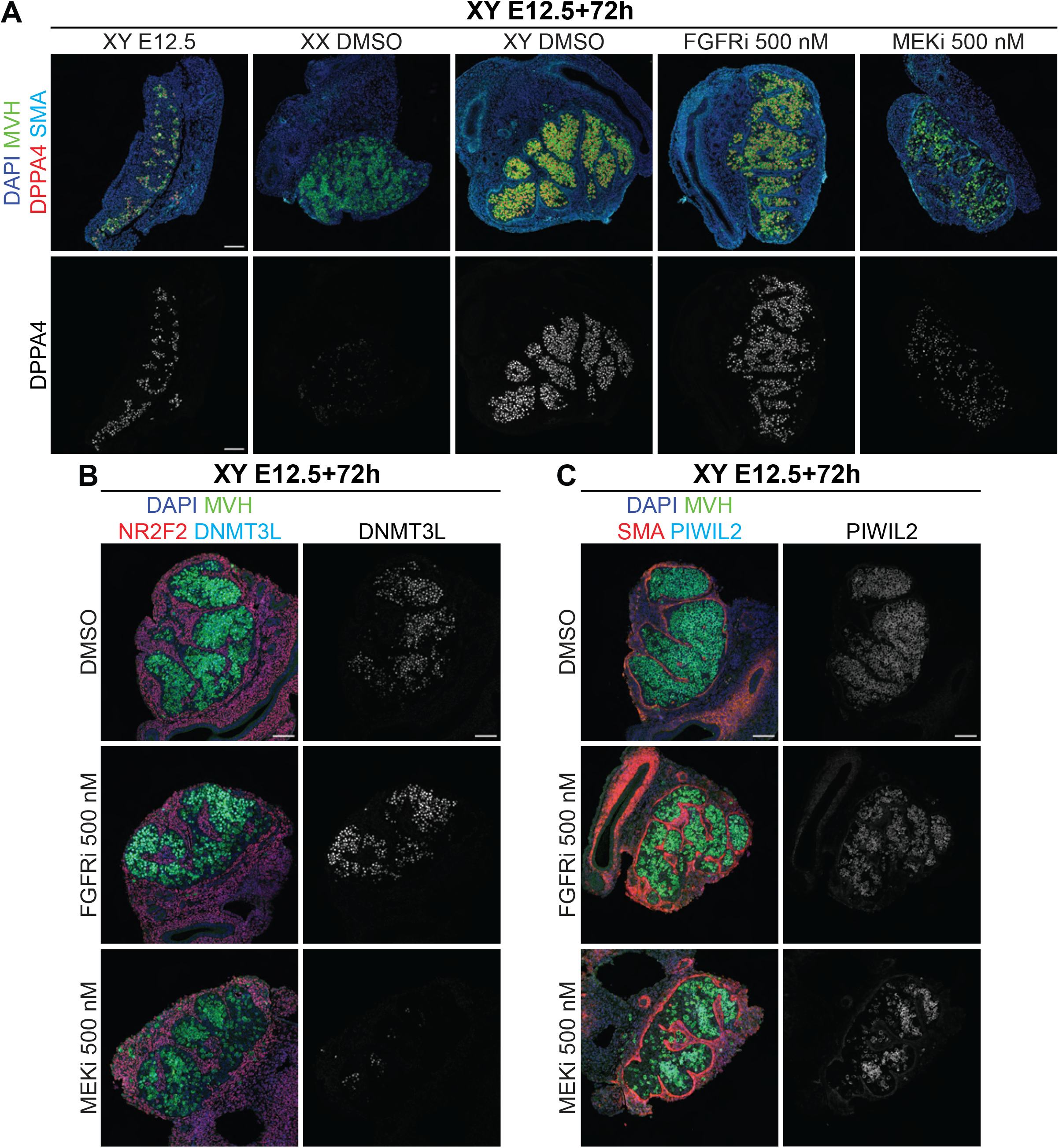
Widefield view of images displayed in Figure 3. Immunofluorescent images of XY E12.5 gonad-mesonephros or XY or XX E12.5 gonad-mesonephros tissue cultured with DMSO or 500nM of FGFRi or MEKi for 72h. **A)** Immunofluorescent images demonstrating DPPA4 localisation. Top panel: DAPI (blue), MVH (green), DPPA4 (red) and SMA (cyan). Bottom panel: DPPA4 (grey). **B)** Immunofluorescent images demonstrating DNMT3L localisation. Left panel: DAPI (blue), MVH (green), NR2F2 (red) and DNMT3L (cyan). Right panel: DNMT3L (grey). **C)** Immunofluorescent images demonstrating PIWIL2 localisation Left panel: DAPI (blue), MVH (green), SMA (red) and PIWIL2 (cyan). Right panel: PIWIL2 (grey). Scale bar: 100μm. Replicates: n=3-4.

**Figure S6:**
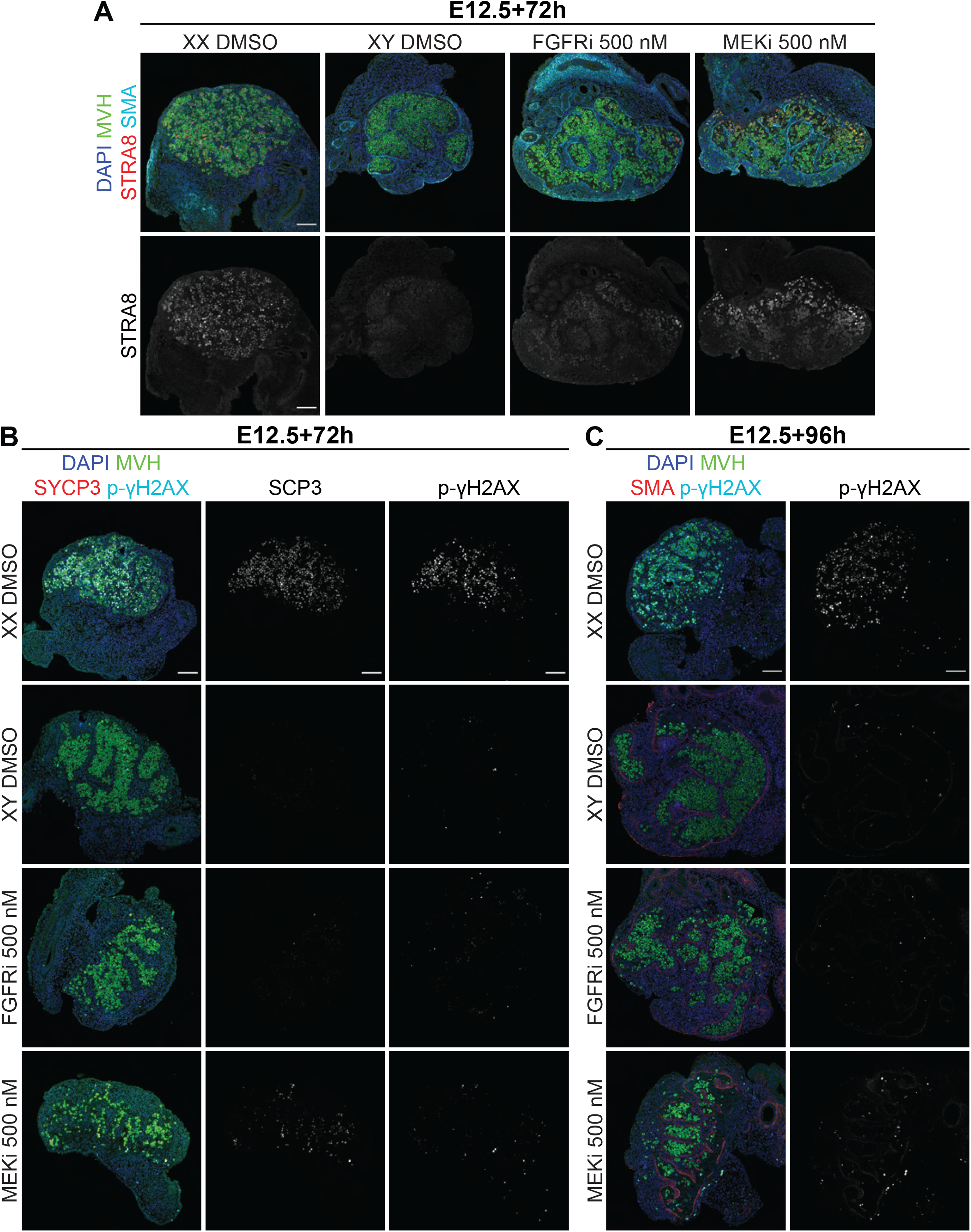
Widefield view of images displayed in Figure 4. Immunofluorescent images of XY or XX E12.5 gonad-mesonephros tissue cultured with DMSO, FGFRi or MEKi for 72h (A-B) or 96h (C). **A)** Immunofluorescent images demonstrating STRA8 localisation. Top panel: DAPI (blue), MVH (green), STRA8 (red) and SMA (cyan). Bottom panel: Stra8 (grey). **B-C)** Immunofluorescent images demonstrating SCP3 and phospho-γH2AX (p-γH2AX) localisation. Left panel: DAPI (blue), MVH (green), SCP3 (red; B) or SMA (red; C) and phospho-γH2AX (cyan). Middle panel: SCP3 (grey; B). Right panel: p-γH2AX (grey). Replicates: n=3-4. Scale bar: 100μm.

**Figure S7:**
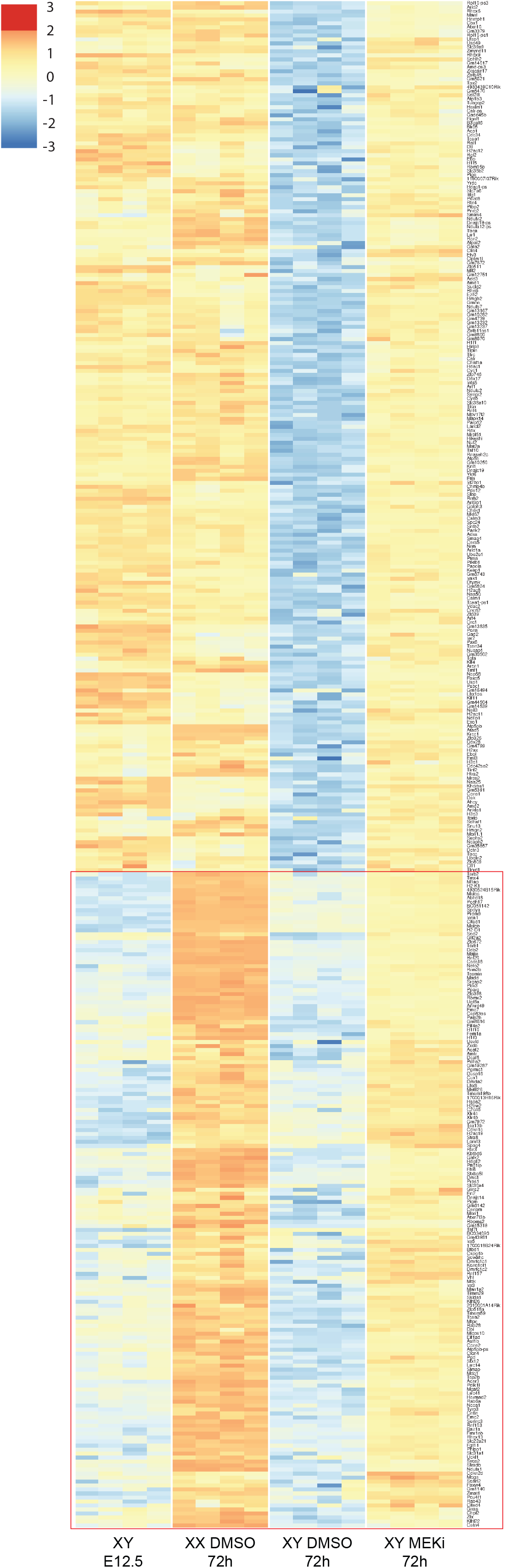
Heatmap of 382 72h MEK1/2 dependent genes expressed higher than expected common in 72h XX germline specific genes. Genes which were expressed higher than expected in XY E12.5+72h MEKi vs XY E12.5+72h DMSO and were present in the 72h XX germline specific genes dataset (identified as genes which were upregulated in XX E12.5+72h DMSO vs XY E12.5+72h DMSO) were assessed. Of these 382 genes, 218 genes were highly expressed in XY E12.5 germ cells and XX E12.5+72h DMSO germ cells and were therefore not considered informative. 164 genes were not or were lowly expressed in XY E12.5 germ cells compared to XX E12.5+72h DMSO germ cells and were therefore considered more reliable female germline differentiation genes (identified by red box). Genes with an FDR <0.05 and |logFC| >0.585 (equivalent to |FC| >1.5) were considered differentially expressed.

**Figure S8:**
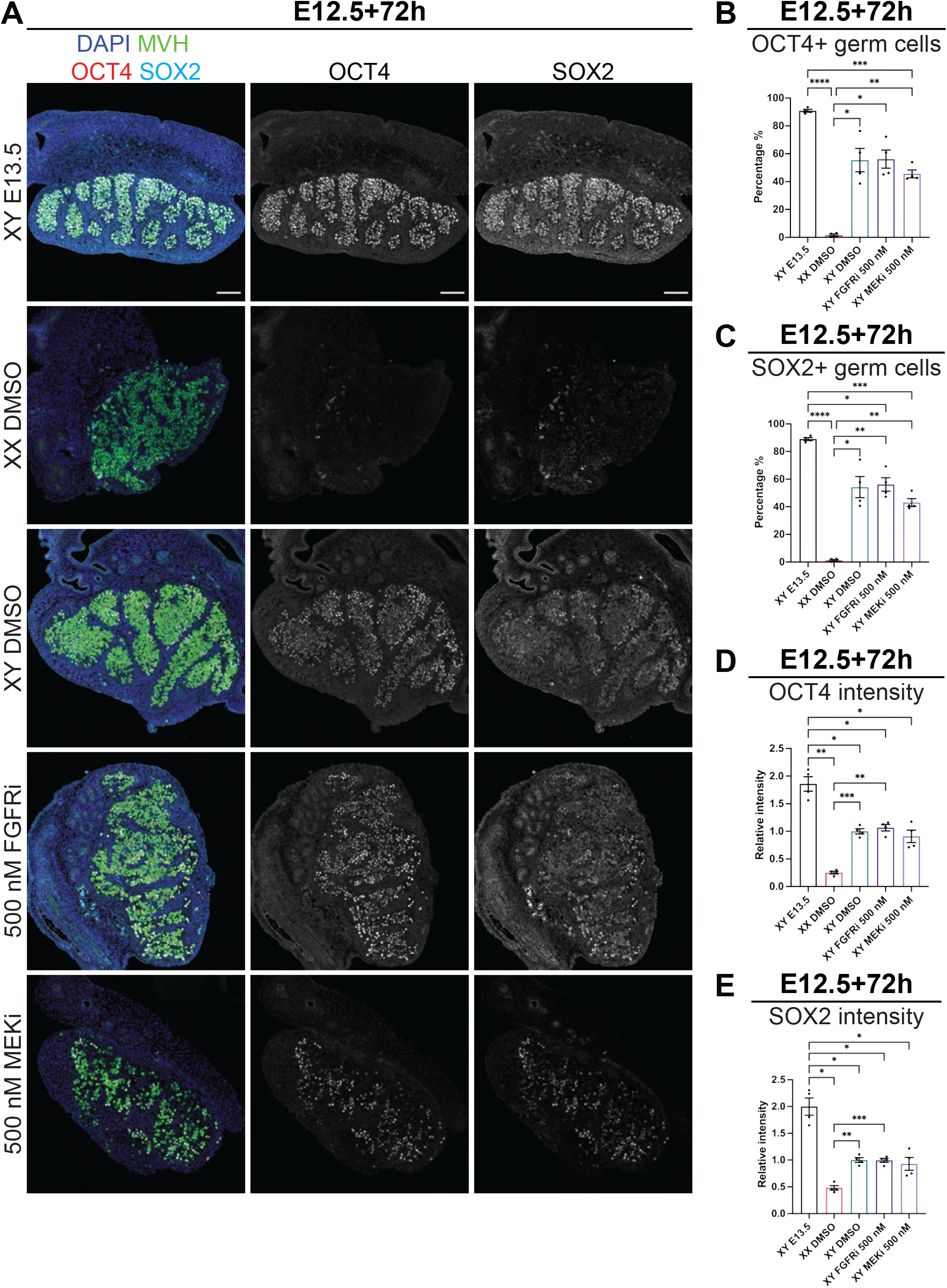
FGF and MEK1/2 inhibition does not result in abnormal maintenance of pluripotency markers. Immunofluorescent analysis of XY E13.5 gonad-mesonephros tissue or XY or XX E12.5 gonad-mesonephros tissue cultured with DMSO or 500nM of FGFRi or MEKi for 72h. **A)** Whole view immunofluorescent images demonstrating OCT4 and SOX2 localisation. Left panel: DAPI (blue), MVH (green), OCT4 (red) and SOX2 (cyan). Middle panel: OCT4 (white). Right panel: SOX2 (white). Scale bar: 100μm. **B-C)** Percentage of OCT4+ (B) or SOX2+ (C) germ cells calculated from immunofluorescent images. **D-E)** OCT4 (C) or SOX2 (E) intensity in germ cells relative to XY DMSO control set at 1.0, calculated from immunofluorescent images. Replicates: n=4. Statistics: Brown-Forsythe and Welch ANOVA with Dunnett’s T3 multiple comparisons. Error bars: mean ± SEM. Significance between controls and treatment: *P < 0.05, **P < 0.01, ***P < 0.001, ****P < 0.0001.

